# Genetic affinities and adaptation of the South West coast populations of India

**DOI:** 10.1101/2022.03.14.484270

**Authors:** Lomous Kumar, Anuhya Chowdhari, Jaison J. Sequeira, Mohammed S. Mustak, Moinak Banerjee, Kumarasamy Thangaraj

## Abstract

Evolutionary event has not only transformed the genetic structure of human populations but also associated with social and cultural transformation. South Asian populations were formed as a result of such evolutionary events of migration and admixture of genetically and culturally distinct groups. Most of the genetic studies pointed to large-scale admixture event between Ancestral North Indian (ANI) and Ancestral South Indian (ASI) groups, also additional layers of recent admixture. In the present study we have analyzed 213 individuals inhabited in South West coast India with traditional warriors and feudal lord status and historically associated with recent migrations events and possible admixture with Indo-Scythians, Saka, Huns and Kushans, whose genetic links are still missing. Analysis of autosomal SNP markers suggests that these groups possibly derived their ancestry from some groups of North West India having additional Middle Eastern genetic component and also their separation history suggests very early separation from North West Indian and Gangetic plain Indo-Europeans during late bronze or Iron age, most probably following central India and Godavari basin to South West coast. Higher distribution of west Eurasian mitochondrial haplogroups also points to admixture through maternal lineage. Selection screen using genome wide genealogy approach revealed genetic signatures related to their long-term coastal food habits. Thus, our study suggests that the South West coastal groups with traditional warriors and feudal lords’ status are of a distinct lineage compared to Dravidian and Gangetic plain Indo-Europeans and are remnants of very early migrations from North West India following Godavari basin to Karnataka and Kerala.

## Introduction

South West coast of India, which includes Konkan and Malabar region, is home to enormous cultural, linguistic and religious diversity emerged from over a millennium of migration, admixture and cultural assimilation and development. In the long history of this region, multiple communities like Greeks, Scytho-Iranians, Kushans, Jewish tribes, Zoroastrians, Moghuls, Muslims, Dutch, Portuguese, English, etc. have left indelible impact on population structure in the form of distinct layers. This highly diverse region also harbors many caste groups linguistically belonging to either Dravidian family (Malayali and Tulu) or Konkani branch of Indo-Aryan language family and historically falling under either priestly status (Havik and Hoysala) or warrior (Nair and Thiyya) and landlord (Bunt) status. Distinct culture, anthropological and historical records relate their origin to ancient migration either from North West India or from the region near the Gangetic plain (Ahichhetra) (Fuller 1976). Ahichhatra is an Iron age archeological site of Painted Grayware Culture (PGW) in Gangetic plains of North India. According to anthropologist, C. J. Fuller, both Nair and Bunts might have a common origin from Ahichhatra. They were brought to the west-coast very early during 375 CE by Kadamba king Mayura Varma along with some groups with Brahminical status like Nambudhiri and Tulu brahmins (Fuller 1976). In the long history of the region different dynasties like Kadamba, Chalukya, Rashtrakuta and Alupa used these groups with warrior status as soldiers. There is a similar kind of mentioning in some historical texts like *Keralolpathi* and Tulunadu *Grama Paddathi*, where they are mentioned as Naga warriors. Thiyya and Ezhava of Malabar also have a separate claim about their warrior status. Although historians believe that Thiyya and Ezhava have migranted from Ceylon (Sri Lanka) bringing palm cultivation and mainly involved in toddy tapping and agricultural works, some written records (Pillai 1970) suggest their origin from Villavars of Chera dynasty and working as soldiers. They are called Thiyya in Malabar while Billavas in Tulunadu. Bunts and Nairs along with some sects of Thiyya and Ezhava practice matriarchy even today. Historical evidence also suggests their contact with populations from the Middle East very early since Kadamba dynasty period and also with Europeans later in the history. Nature of these contacts were majorly commercial but had an impact on the society through the spread of religions like Judaism, Islam and Christianity.

Most of the previous studies related to migration and admixture history of India were mainly focused on major migration of R1a carrying populations in Bronze age Steppe which later were prevalent as ANI component among groups recognized with priestly status in caste hierarchy (Narasimhan, et al. 2019). Before the arrival of this component, populations of South Asia mainly comprised of Ancestral South Indian (ASI) component, which was prevalent among bronze age Indus valley civilization people. But at the verge of fall of Indus civilization around 4000 years ago (Dutt, et al. 2018) both these components admixed and gave rise to an ANI-ASI cline rooting all the castes and tribal groups of the Indian subcontinent. However, our earlier studies, which tried to model these populations as simple admixture of two major components (ANI and ASI) in varying proportions failed mostly in case of caste groups of both North and South India (Moorjani, et al. 2013; Narasimhan, et al. 2019). Reason we proposed behind this was the complex nature of admixture history in the form of multiple layers of ancestries. More specifically in South West coast of India, studies with some unique population groups like Jews, Parsees and Roman Catholics have already pointed about the additional layers of admixture among them (Chaubey, et al. 2016; Chaubey, et al. 2017; Kumar, et al. 2021). Previous genetic studies based on Y chromosomal microsatellite markers also found more west Eurasian or North West Indian genetic influence in gene pool of Nairs and Ezhava (Nair, et al. 2011; Mahal and Matsoukas 2018). While another genetic study based on the human leukocyte antigen (HLA)-A, -B and -C diversity found greater Dravidian influence along with traces of admixture with west Eurasian, Mediterranean, Central Asian and East Asian populations (Thomas, et al. 2006). Detailed genetic history of these groups with either warrior or feudal lord status having footprints of later migrations to South West coast was majorly uncovered. Therefore, in our present study we included population groups of non-Brahminical, warrior and feudal lords’ status like Nair, Thiyya and Ezhava from Kerala and Bunt and Hoysala from Karnataka from South West coast of India in order to dissect history of their genetic origin and adaptation, as well as to unravel further layers of ancestral component manifesting as a result of admixture of Bronze age, Iron age or later migrations.

## Results

### Distinct clustering of South West coastal populations in the context of all Eurasian populations

We first performed principal component analysis (PCA) to gain insight into population structure. Interestingly, we found that south Asian populations are distributed into five major clusters along the ANI-ASI cline, with groups from Pakistan/North West India at one extreme; while many Dravidian tribes at the other extreme (Fig1b). Most of the Gangetic plain Indo-Europeans along with a few North West Indian individuals formed third major cluster, while fourth major cluster comprised of heterogenous groups, including Indo-Europeans from north India, groups with priestly status from Konkan, our five study groups (Nair, Thiyya, Bunt, Ezhava and Hoysala) and some of the Dravidian groups from geographical vicinity of our study groups. In this cluster, Nair and Hoysala are near Havik and Karnataka brahmins much closer to third cluster of Gangetic plain Indo Europeans. Bunt is adjacent to this group but further away in the cline. Thiyya individuals are found both along with Nairs but fewer and also majority are further away towards Dravidian groups of this cluster along with Karnataka Gaud. Ezhava are last in this cluster with Reddy caste and some of the Dravidian populations, such as Kuruba and Kunabi. We also found an interesting displacement of a few Sikh_Jatt and Muslim_Kashmiri individuals towards this fourth heterogenous cluster of our study groups (Nair, Bunt, Thiyya, Ezhava and Hoysala) (Fig 1b).

**Fig 1.**
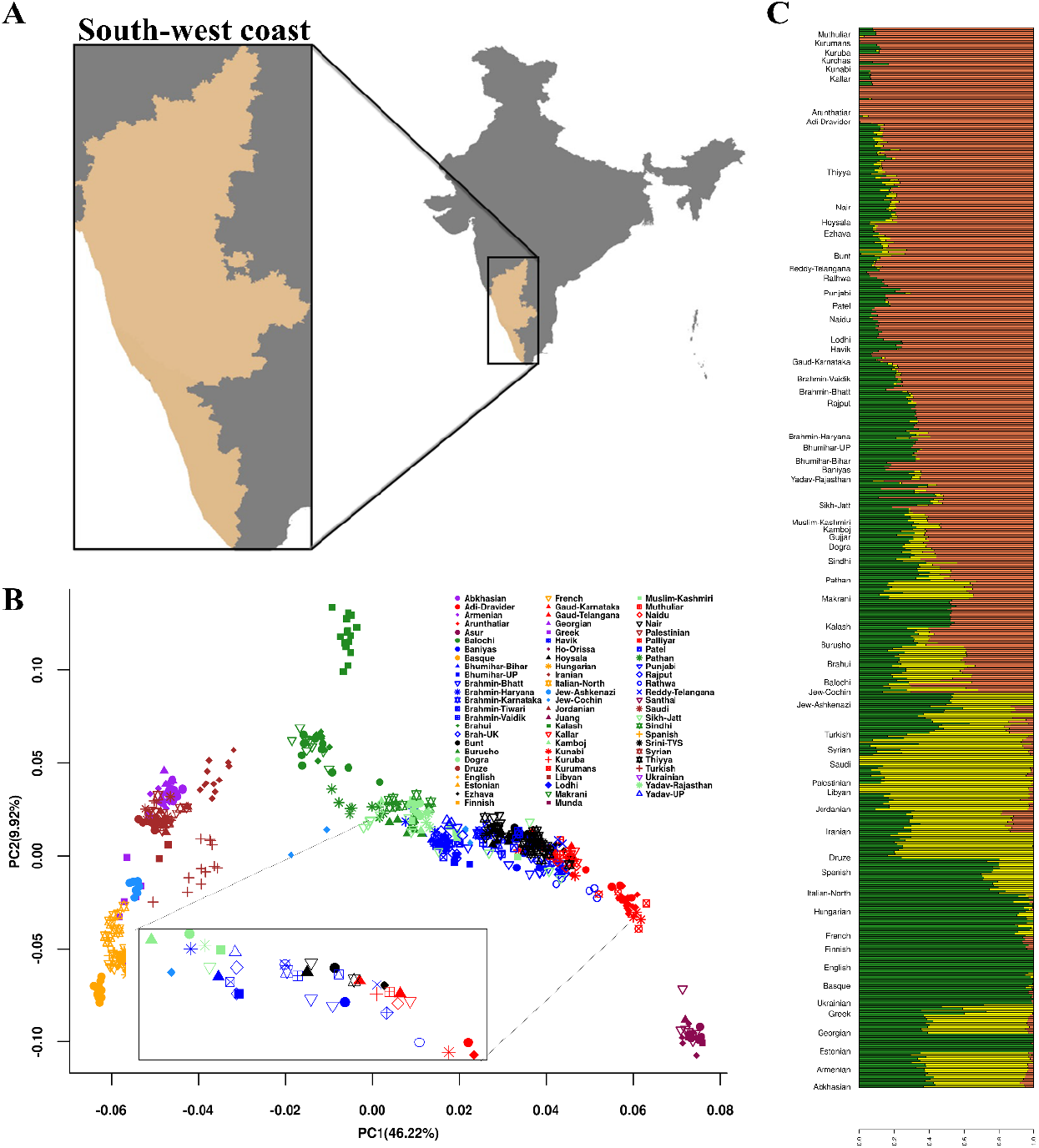
Sampled region, PCA and Admixture plots. **A.** Map of India with regions of Konkan and Malabar coast from state of Karnataka and Kerala respectively (shaded with yellow color representing South West coast), inset image shows South West coast of Konkan and Malabar from Karnataka and Kerala. **B.** Biplot of Principal Component Analysis (PCA) of South West coastal groups with modern Eurasian populations with first two components. Inset is biplot of the population mean of first two principal components. **C.** Stacked bar plot of the ADMIXTURE analysis with K=3, using modern west Eurasian populations as reference. Populations are arranged from bottom to top.

In order to infer the ancestral genetic components in the context of modern Eurasians and to further enquire about the clustering pattern found in PC analysis, we used model-based approach in ADMIXTURE (Alexander, et al. 2009). Surprisingly, population groups placed in fourth cluster of our PC analysis, except Dravidian groups like Kuruba and Kunabi, showed additional prominence of yellow component, which is characteristics of populations from Middle East and present in significant proportions among populations from Pakistan (Pathan, Balochi and Makrani) and North West India (Kamboj, Gujjar, Muslim_Kashmiri, Dogra and Yadav_Rajasthan) (Fig 1c). While other populations in the ANI-ASI cline, such as Gangetic plain Indo-Europeans (Brahmin_Tiwari, Bhumihar_Bihar, Patel, Lodhi etc.) and Dravidian cates and tribe groups are lacking this, with exceptions of Kuruba and Kunabi populations.

We further tested for the admixture history and allele sharing pattern in the five study groups of South West coast by utilizing admixture F3 and Dstatistics methods in qp3Pop and qpDstat tools of ADMIXTOOLS 2 package in R. Admixture F3 was run in the form F3 (X, Palliyar; Nair/Bunt/Thiyya/Ezhava/Hoysala) using Palliyar as proxy for ASI source of ancestry and population X as different West Eurasians and south Asian populations. We found that Nair and Thiyya shows significant F3 statistics with Middle Eastern population (Iranian, Druze) in contrary to populations from either Caucasus or Europe, which is a characteristic of most of the groups with Indo-European affinity (Fig S1a-b). Although populations like Bunt, Hoysala and Ezhava show highest F3 statistics with either Caucasus or European populations, but groups from Middle East rank third or fourth in terms of higher F3 statistics (Iranian, Druze) (Fig S1c-e). However, none of the five populations (Nair, Bunt, Thiyya, Ezhava and Hysala) showed significant admixture F3 statistics with each other.

We then calculated Dstatistics in the form F4 (pop1, pop2, Steppe/Yamnaya, Yoruba) and F4 (pop1, pop2, Iran_N, Yoruba) to compare relative affinity of various modern Indian populations for Steppe vs Iranian ancestry in comparison to our study groups. Here, pop1 is various modern Indian cline groups and pop2 is our study group (Nair/Bunt/Thiyya/Ezhava/Hoysala) of the South West coastal India.

In the scatterplot, Nair shows higher Steppe and Iran_N affinity outcompeting all other south Asian groups except groups from Pakistan, north west India and few populations like Bhumihar_Bihar and Rajput from Haryana (Fig S2a). Interestingly, Nair displayed comparatively more gene flow from Iran_N than Cochin Jews and also other populations of South West coast or Godavari basin (Reddy and Vaidik Brahmins). Bunt and Hoysala exhibit similar trend as Nairs (Fig S2c, Fig S2e) but Thiyya and Ezhava show comparatively more shifting of other Indo-European groups towards right side of scatter, so more groups are outcompeting Thiyya and Ezhava in terms of geneflow from both sources viz. Steppe and Iran_N (Fig S2b, Fig S2d).

In the Maximum likelihood tree constructed using TreeMix v.1.12 (Pickrell and Pritchard 2012), placement of all five groups was consistent with their clustering in PCA analysis, with Nair and Hoysala among North Indian Indo-European caste groups, and Thiyya and Ezhava are more towards Dravidian cluster (Fig S3). however, we didn’t observe any population-specific drift among the sample groups. (Fig S3).

### Fine scale population structure using haplotype-based approach

To gain better understanding of population structure and haplotype sharing pattern of five population groups of South West coast with modern Eurasians, we used haplotype-based approach with ChromoPainter (Lawson, et al. 2012) and fineSTRUCTURE (Lawson, et al. 2012). fineSTRUCTURE (Lawson, et al. 2012) clustering divided all Indian samples into two major clades, one with North west Indian and Gangetic plain groups and other clade with south Indian groups. South Indian clade was further divided into two major subclades, where one of them keeps Bunt and Hoysala individuals together, while the other one is heterogenous clade comprising of all remaining populations in different branches together (Fig S4a-b). This cluster includes Nair, Thiyya, Ezhava, some individuals of Bunt and Hoysala, Godavari basin populations such as; Reddy, Vaidik brahmins and Naidu and also Dravidian groups namely; Kuruba, Kunabi and Kurchas from Karnataka and Kerala. Havik and Karnataka brahmins are also in this cluster along with Hoysala, Bunt, Nair and Thiyya (Fig S4a-b).

### Ancient ancestral contribution to South West coastal groups

We applied admixture modelling approach with two different sources of Iranian ancestry viz. pre-Bronze age Namazga_CA and Bronze age Indus_Periphery group using qpAdm of ADMIXTOOLS 2 to compare the ancient contributions into ancestry of five groups of South West coast and other south Asian populations. In the first approach, we used Andamanese Hunter-Gatherers (AHG), Namazga_CA and Steppe_MLBA as left groups. Among all south Asian groups tested Namazga_CA component was comparatively higher in proportion in North West Indian groups, such as Gujjar (0.53), Kamboj (0.46) and Pathan (0.45) population from Pakistan. Surprisingly, all our five sample groups have higher proportion of Namazga_CA ancestry (Fig 3b) (Supplementary Table S7) along with other North West Indian groups, such as Muslim_Kashmiri, Dogra and Yadav_Rajasthan in comparison to Gangetic plain populations like Brahmin_Tiwari (0.34), Bhumihar_Bihar (0.35), Srivastava (0.35) etc. and also other Dravidians like Mala (0.32), Naidu (0.39), Palliyar (0.25) and Ulladan (0.24) from south India.

In the admixture modelling with Bronze age sources (AHG, Indus_Periphery and Western/Central Steppe MLBA), we found that after Gujjar (0.71) and Kamboj (0.63), Nairs (0.61) are the group with higher Indus periphery component from Bronze age (Fig 3a) (Supplementary Table S8). Bunt is with similar contribution from Indus periphery group (0.44) and also Thiyya, Ezhava and Hoysala along with groups from North West India like Yadav_Rajasthan, Dogra and Muslim_Kashmiri are higher in distribution of Indus periphery component than most of the Gangetic Plain populations and other Dravidian population in south India (Fig 3a) (Supplementary Table S8).

We further tested all five groups from South West coast to fit in the Admixture graph topology comprising both modern and ancient population groups, using qpGraph function in ADMIXTOOLS 2. We tested different alternate topologies for all five groups to arrive at best fitted model with maximum likelihood score (closer to zero) and here we are showing only those results having best fit. For Nairs, we obtained a graph topology with best fit (likelihood score 2.94125), showing a pattern of admixture typical of ANI-ASI admixture from an ASI group similar to Palliyar and an ancient ghost population ANI formed by admixture between Indus population and Yamnaya like Steppe group (Fig S5a). In addition to this simple ANI-ASI admixture, Nairs also require another source group for Middle East ancestry from Bactria-Margiana-Archeological-Complex (BMAC).

We used Brahmin_Tiwari population from Uttar Pradesh to represent the Gangetic Plain group, which was in recent study found to be highest in carrying Steppe component (Narasimhan, et al. 2019). While Nair required additional Middle Eastern component, Brahmin Tiwari from Gangetic plain was best fitted without it in a simple ANI-ASI admixture model (Fig S5a). We applied admixture modelling approach with other study groups and we found that all of these groups from South West coast India are best fitted in the same model of additional Middle Eastern component along with ANI-ASI admixture required by other Indo-Europeans and Dravidian castes and tribes (Fig S5b-e). Likelihood scores for Thiyya (3.294107), Bunt (2.919086), Ezhava (2.817161) and Hoysala (1.7) were maximum and closer to Zero value.

We also estimated the Admixture graph model for one of the North west Indian group to compare them with Gangetic plain populations and our South West coastal study population. In this case, we used Gujjar population which was showing highest proportion of Iranian ancestry in admixture modelling with source groups Namazga_CA (0.53) and Indus Periphery group (0.71) compared to all other south Asian populations.

In admixture graph modelling, Gujjar population also was best fitted in a model, where they require additional Middle Eastern component (Fig S5f) along with the basic model of ANI-ASI admixture (Moorjani, et al. 2013). The likelihood value for this admixture model for Gujjar population (2.3073) was maximum and very closer to zero (Fig S5f).

### Estimation of Effective Migration Surface

Effective migration surface is the visual representation of geographical population structure in terms of effective migration. This representation of population structure highlights potential regions of higher-than-average and lower-than-average historic gene flow (Petkova, et al. 2016). We applied this method to our study groups from South West coast of India along with reference populations on ANI-ASI cline. We included in this analysis mainly the population groups from Indo-European and Dravidian linguistic family and excluded groups from Tibeto-Burman and Austroasiatic linguistic family, since we were mainly interested in migration events related to ANI-ASI admixture.

This method uses pairwise genetic dissimilarity matrix calculated from genotype data and geographical coordinates of samples as raw input and derive the posterior distribution of effective migration and diversity rates using MCMC iterations. Here, we first used varying number of Demes viz. 150, 175, 200, 225 and 250 demes five times each to run MCMC iteration and found 200 demes as best fitted for combined sample set. We further proceeded with main MCMC algorithm using 200 demes and varying acceptance proportions for proposal distributions to arrive at best suitable proposal distributions. After this, by applying these conditions, we ran MCMC algorithm for 5 million MCMC iterations, 1 million burn-in and 10000 sample iterations. Based on final posterior distribution of effective migration and diversity rate, we plotted the migration surface.

We observed that there are five distinct regions across India having higher than average migration rate (Fig S6a). First region is in the North India (in Jammu & Kashmir) and is continuous with North west India. Second region is mid-Gangetic plain of Uttar-Pradesh and Bihar. Third region is in central India, which is continuous with some regions in further west and also north. Fourth and fifth regions are in the Godavari basin and Karnataka respectively, which are almost continuous to each other and some nearby regions extending further to coastal Karnataka. This reflects continuity in gene flow pattern across this large region in south India.

We also observed many regions of lower-than-average migration rates. One such region is in North West India separating two regions of higher-than-average migration rates in North India (region 1) and Central India (region3). This region makes clear boundary or obstacle in gene flow between these two regions (Fig S6a).

Other such regions of higher migration rates are between mid Gangetic plain and Godavari basin region and also between Karnataka and Central Indian region. Southernmost part of India has another region of lower-than-average migration rates along with minor continuous regions of east coastal India. Fig S6b represents the convergence of main MCMC algorithm, while Fig S6c represents observed vs fitted dissimilarities between demes.

### Temporal dynamics of Effective Migration rates

We further explored the geographical gene flow pattern or migration rates dynamics over passage of time. Using matrix of similarity of shared long segments of haplotypes, also referred as Long Pairwise Coalescent Segments (lPSC), among individuals of populations and varying lengths of these shared segments gives time dependent estimates of effective population size and migration rates, a method implemented in MAPS (Al-Asadi, et al. 2019). Further, using geographical coordinates of samples along with genetic data, we inferred the geospatial patterns of migration rates and population size changes with time.

For this purpose, we obtained the phase genotype data and IBD segments using Beagle5.1 (Browning, et al. 2018) and Refined-IBD (Browning and Browning 2013) tools, then using predefined pipeline we derived the matrix of IBD sharing among individuals. We used three different lPSC segment, lengths windows viz. 1-5 cM and 5-10 cM which corresponds to genetic time frame of 90 generation and 22 generations ago from present. We used IBD sharing matrix and geographical coordinates of samples and obtained the posterior distribution of the parameters effective migration rates and effective population size using MCMC algorithm. Using 200 demes, we first obtained the best fitted variance proposals so that all values lie between 10-40 % of range. Then, we ran the main algorithm 10 times using 5 million MCMC, 1 million Burn-in and 10000 sample iteration and inferred the posterior distribution of parameters of effective migration rates and population sizes.

#### Effective migration surface in 1-5 cM PSC length window

In the PSC length segment range of 1-5 cM, which corresponds to mean coalescence time of 90 generation ago, we found that the North West India is having higher than average distribution of migration rate (Fig 2a). Other regions include central India and upper Godavari basin and these two regions of higher-than-average migration rates are almost continuous to each other without any boundary. Another region of very high migration rate is near south-east coast India, but this region is separated from Godavari region by a boundary of lower-than-average migration rate (Fig 2a).

**Fig 2.**
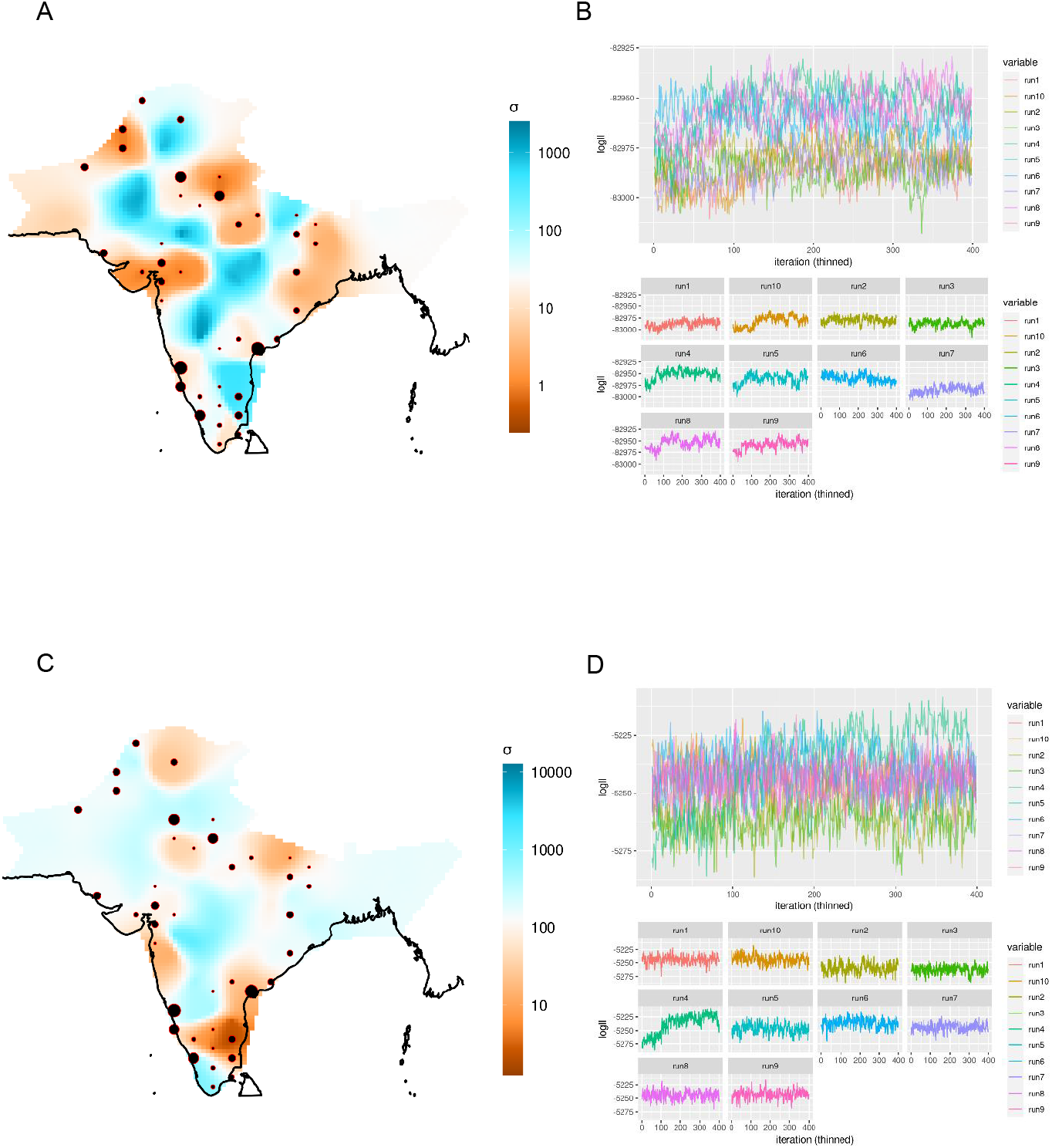
Spatial pattern of effective migration rates through time. **A**. Contour plot of effective migration rate in 1-5cM lPSC length corresponding to a timeframe of 90 generations ago. Blue colour represents regions of higher-than-average migration rates, while orange colour shows regions of lower-than-average migration rates and **B.** corresponding Markov Chain Monte Carlo iteration chain convergence. Plot shows the log likelihood distributions in three individual MCMC runs. **C.** Contour plot of effective migration rate in 5-10cM lPSC length corresponding to a timeframe of 22 generations ago and **D.** corresponding MCMC iteration chain convergence in three individual runs.

**Fig 3.**
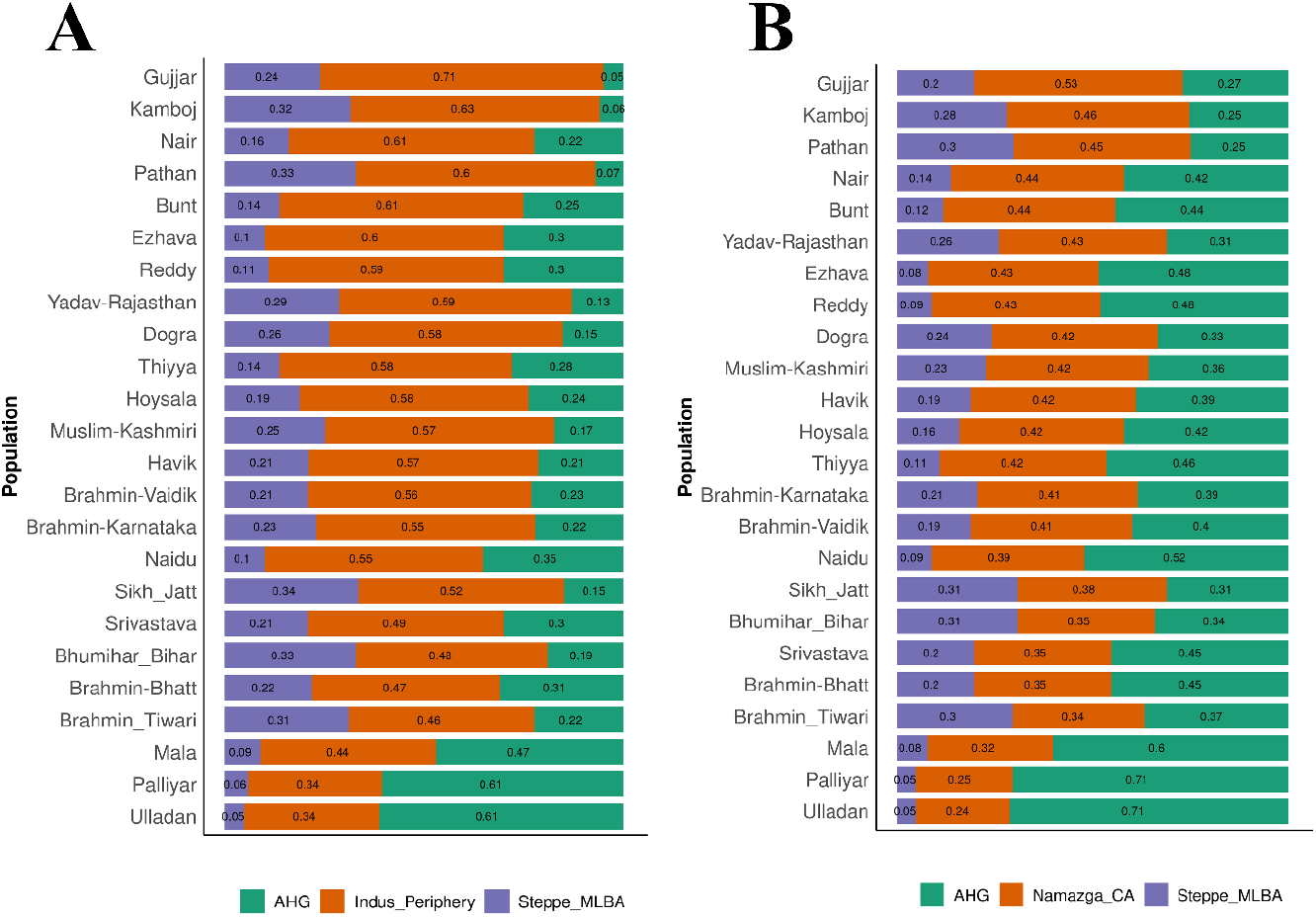
Admixture modelling to infer contribution of ancient ancestral west Eurasian source populations. **A.** Admixture modelling of South West coastal groups along with some other modern Indian populations using Andamanese Hunter Gatherers (AHG), Indus Periphery group and Middle and Late Bronze age Steppe groups (Steppe_MLBA) as ancient sources. Each color in bar plot represents fraction of ancestry from individual ancient source groups. **B.** Admixture modelling of South West coastal groups using Andamanese Hunter Gatherers (AHG), Namazga Chalcolithic group and Middle and Late Bronze age Steppe groups (Steppe_MLBA) as ancient sources.

#### Effective migration surface in 5-10 cM PSC length window

The PSC length window of 5-10cM corresponds to a mean coalescence time of 22 generations ago from present. The migration surface at this time shows a diffused pattern in the regions of higher-than-average migration rates, like North West India, central India and Godavari basin, while regions in Gangetic plain have now lower than average migration rates (Fig 2c). Upper Gadavari basin is still a region of high migration rate compared to all other parts of India. Regions of south-east coast, which were earlier regions of very high migration rates now shows different pattern and becomes a region of very low migration rates. An additional region of higher-than-average migration rates appears in southern most part of India, which is secluded by a boundary of low migration rates (Fig 2c). Regions of North India near Jammu and Kashmir have now lower than average migration rates along with one more such region near north west India. Parts of central India, Godavari basin and parts of Karnataka including coastal Karnataka are now with similar migration rates (i.e., higher than average migration rates).

### Population separation history and effective population size

We further explored the population separation history of South West coast population with populations from North West India, Gangetic plains, Middle East, European and African population using a genome wide genealogy-based approach in Relate (Speidel, et al. 2019). First estimating the branch lengths of genealogy tree and then coalescence rate between pair of haplotypes and applying MCMC algorithm, we estimated the effective population size.

As it is shown in the stairway plot (Fig S7), first separation of all the groups occurred from African population much after 100 KYA followed by Bronze age separation of Indian populations from Middle East and European groups after 5 KYA. Also, as evident from the stairway plot, separation of South West coastal groups started much earlier, before the timeframe of 1000 years ago (may be late Bronze age or Iron age separation).

### Genome wide genealogy-based screening for positive selection

Using genome wide genealogy-based approach in Relate (Speidel, et al. 2019), we performed screening for possible candidate regions for positive selection. This approach is based on the rapid spread of favorable mutations under positive selection in a particular population. This approach also called Relate selection test is based on estimation of speed of spread of favorable allele compared to a competing allele over a particular time frame. By calculation of p-value of positive selection using above approach, we identified 7 strong signals of positive selection genome wide (p-val < 10^-5^) (Table 1). Of the five major regions we have listed in Table-S1c, *WDR5* involved in skeletal system development and megakaryocyte differentiation. We also found two regions of strong selection in genes *TPK1* and *DMRT1* also previously reported in earlier selection screening (Kimura, et al. 2007; Liu, et al. 2013; Schaschl and Wallner 2020). *TPK1* is involved in Thiamine metabolism and may be involved in adaptation to Thiamine rich diet in the form of more fish and other seafoods.

**Table 1.** Signal of positive selection inferred by Relate selection screening using genome wide genealogy

| ID | Lower age | Upper age | Ancestral/Derived | DAF | P value | Gene | Function |
| --- | --- | --- | --- | --- | --- | --- | --- |
| **rs34856382** | 5211.3 | 10850.7 | T/C | 3 | -1.59478E-05 | *WDR5* | Skeletal system development and Megakaryocytes differentiation |
| **rs12264666** | 3494.7 | 7542.2 | A/G | 3 | -1.59478E-05 | *KIAA1217* | Skeletal system development |
| **rs77214450** | 2378.2 | 6970.7 | G/A | 4 | -9.31754E-05 | *PALLD,CBR4* | Keratinocyte development |
| **rs6957904** | 3541.6 | 8904.7 | A/G | 4 | -9.31754E-05 | *TPK1,OR2A1, OR2A2, OR2A7* | Thiamine metabolism |
| **rs364477** | 3430.3 | 6884.5 | T/C | 4 | -9.31754E-05 | *DMRT1,DMRT2, DMRT3* | Male fertility |

*DMRT1* is male fertility related gene. Two more interesting signals were found in *PALLD* and *CBR4* gene involved in keratinocyte development and lipid metabolism, respectively. Both these regions were earlier reported as signal of positive selection among Europeans based on iHS, LRH and FST based approach (Lappalainen, et al. 2010).

As clearly evident from the marginal phylogenetic tree around the SNP rs6957904 (Fig S8a) in the gene *TPK1*, the SNP rs364477 (Fig S8b) in the gene *DMRT1*, the SNP rs77214450 (Fig S8c) in the gene *PALLD and* the SNP rs12264666 (Fig S8d) in the gene *KIAA1217* the spread of ancestral allele is more among all other populations compared to African population. This spread is more in French, Iran, North West India, Gangetic plain groups and South West coast populations. Among these, later groups extent of ancestral allele spread is comparatively more among North West Indian and South West coastal populations than among Gangetic plain population in India.

### Mitochondrial haplogroup distribution among South West coastal groups

We compared the mtDNA haplogroup diversity among South West coastal groups and found that maternal lineage is very diverse among Nair, Thiyya and Bunt of Konkan and Malabar coast (Fig S9a) (Supplementary Table S1). We observed prevalence of five major haplogroups (M, R, U, N and H), of these haplogroup M is the most prevalent followed by U and R. Surprisingly we observed higher proportion of haplogroup H among Bunts of Konkan coast and also with lesser frequency among Nair and Thiyya from Malabar (Fig S9a) (Supplementary Table S1). Haplogroup N was present in significant fraction of Thiyya population, while haplogroup U was with highest occurrence among Nairs.

Among the subgroups of haplogroup M, sub haplogroup M3, M35 and M5 was present with highest frequency among Nair (M3 = ~0.14), Thiyya (M35= ~0.12) and Bunt (M5 = ~0.12) (Fig S9b) (Supplementary Table S1). Oldest sub haplogroup M2 was observed with noticeable frequency in all the three groups, while haplogroups M39 and M81 was only present in Bunt (M39 = 0.06; M81 = 0.01) and Thiyya (M39 = 0.01; M81 = 0.01). Some haplogroups like M57a and M64 (earlier observed among linguistic isolate Nihali population) are only observed in Thiyya while haplogroups M30, M61, M65b and M80 were present only among Bunts. Among the subgroups of haplogroup R, haplogroups R6 and R7a was present in significant frequency among Nair (R6 = ~0.08; R7a = ~0.04) and Thiyya (R6 = ~0.05; R7a = ~0.03) while basal group R was found with frequency of 0.02 and 0.067 in Nair and Thiyya, respectively (Fig S9d) (Supplementary Table S1). Haplogroup R30 was present among Nair and Bunt, R5a among Bunt and Thiyya while R8 was found exclusively among Thiyya. Among the subgroups of U haplogroup, U7a, U1a and U2c were found among all three groups, while U2a, U2b and U7b were present only in Nair and Thiyya. Haplogroup U5a was observed only in Bunt population while U9a was found among Bunt and Thiyya (Fig S9c) (Supplementary Table S1).

## Discussion

South West coastal India is a region of high population diversity and with complex genetic history. Some earlier studies with groups like Jews (Behar, et al. 2008; Behar, et al. 2010; Chaubey, et al. 2016), Parsees (Chaubey, et al. 2017) and Roman catholic (Kumar, et al. 2021) already pointed out the complex genetic history and multilayered genetic structure of populations in this region. Also, earlier studies (Moorjani, et al. 2013; Narasimhan, et al. 2019) indeed suggested about the multi layered admixture in the Indian subcontinent specially among caste groups. Therefore, in our current study, we further explored these addition layers of genetic admixture and migrations in caste groups (majorly traditional warriors and landlords) of South West coastal India.

Historical records propose two distinct hypotheses regarding the origin of South West coastal groups of traditional warriors or landlords’ status. According to oral tradition and some earlier genetic studies (Nair, et al. 2011; Mahal and Matsoukas 2018), Nairs and Ezhava have common origin with north west Indian populations, particularly Sikh_Jat, which in turn were historically related to Indo-Scythian tribes (Mahil 1955; Dhillon 1994; Nijjar 2008; Marshall 2013). While some historians relate their origin to Iron age populations of Ahichhatra, Uttar Pradesh (Fuller 1976). The outcome of our PCA analysis with autosomal dataset initially supports both the hypothesis, as populations like Nairs, Bunts and Hoysala from South West coast are closely placed near north and north west Indians. Although Placement of Thiyya and Ezhava more away towards Dravidian clusters points to higher level of local admixture in them. However, clear presence of additional Middle Eastern component (highly prominent among groups that includes Balochi, Pathan and Sindhi from Pakistan and Kamboj, Dogra, Gujjar and Yadav_Rajasthan from North West India) in Admixture analysis and also in our admixture modelling approaches contradicts the hypothesis of Ahichhatra origin of South West coastal groups, while strengthens the hypothesis of their origin from a group closely related to north west Indian Indo-Europeans. Our Admixture F3 statistics also supports this view, as groups like Nair and Thiyya have highest affinity with Middle Eastern population rather than populations from Caucasus and Europe. Although populations like Ezhava from Malabar, Reddy and Vaidik Brahmin from Godavari basin and Gaud from Karnataka also have this Middle Eastern component but in lesser proportions. These groups possibly derived this additional component from ancestral groups of South West coastal population while migration through Godavari basin and Karnataka, as our geospatial and temporal population structure analysis using EEMS and MAPS suggests these regions along with Narmada basin to be key transition zone of gene flow from northwest/north India to south India. Another plausible explanation here can be shared origin from a common ancestral lineage of groups like Reddy from Godavari basin and Gaud from Karnataka with that of the populations of South West coast. None of the group showed significant F3 statistics with Gangetic plain Indo-Europeans, but it was significant with north western groups such as Kamboj, Gujjar and Yadav_Rajasthan only in case of Nair, Bunt and Thiyya. Admixture graph modelling approach also supports the presence of additional Middle Eastern component in comparison to Gangetic plain populations like Brahmin_Tiwari. Same graph model was also applicable for north west Indian group like Gujjar. These results further strengthens the hypothesis of common origin of Nair, Bunt and Thiyya from a population related to those of modern North West Indian groups but definitely not from Indo-Europeans of Gangetic plain or the region near Ahichhetra (Uttar Pradesh) (Fuller 1976).

Our Chromopainter-FineSTRUCTURE analysis clearly suggests that Bunt and Hoyasla share more haplotypes and there was recent admixture between these two groups, while Nair, Thiyya, Ezhava and remaining individuals from Bunt and Hoysala share haplotypes with most of the groups from South West coast like Havik, Brahmin_Karnataka, Kuruba, Kunabi and Kurchas and also Godavari basin like Vaidik Brahmin, Reddy and Naidu, reflecting long admixture history with these groups or groups related to them.

Population separation history of South West coastal group from those of the North West India and Gangetic plain populations using genome wide genealogy-based approach with Relate (Speidel, et al. 2019) clearly indicates separation of these groups from each other long ago (> 2000 years). Probably this is the main region behind presence of large gap in the PCA clusters of South West coast populations and those of north west/north Indian groups. This long-term separation from north west/north Indian populations combined with local admixture to large extent is reflected in more haplotype sharing of South West coast populations with their surrounding caste and tribe groups. The allele spread patterns in the regions of positively selected loci like *TPK1, DMRT1* in our genome wide genealogy-based approach of Relate (Speidel, et al. 2019) depicts further similarity of South West coast population with that of the populations of north west India.

In terms of mitochondrial lineage spread, South West coastal populations exhibit very high diversity with presence of different sub lineages of macro haplogroups M, R, U and also presence of west Eurasian haplogroups HV and H. Although sub haplogroups M, R and U were observed in earlier studies with Indian populations (Kivisild, et al. 1999; Cordaux, et al. 2003; Metspalu, et al. 2004; Palanichamy, et al. 2004; Chandrasekar, et al. 2009), H was observed with very low frequency among the caste groups of south India (Kivisild, et al. 1999; Palanichamy, et al. 2004). We observed high frequency of H haplogroup in Bunt population and also HV in Thiyya population. Presence of high maternal haplogroup diversity among these groups points to possible admixture of diverse maternal lineage in them, which may be true because of their history of practicing of matriarchy.

To sum up, we found a distinct group of populations from South West coastal India, who belong to traditional warriors or feudal lords’ status and have genomic signature of additional Middle Eastern component compared to Gangetic plain Indo-Europeans and other Dravidian caste and tribes. This signature is also typical of some north west Indian populations like Gujjar, Kamboj and Yadav-Rajasthan. PCA clustering and haplotype sharing pattern indicates early population separation and isolation of South West coastal populations from other Indo-Europeans of north and North West India. Study of geographical population structure and time dependent migration rates suggest region of central India (Narmada basin) and Godavari basin to be a transition zone for gene flow from North West or north India to South West India. Population separation history points to much earlier separation of South West coastal groups from North West Indian and high mitochondrial haplogroup diversity along with comparatively higher prevalence of west Eurasian mtDNA haplogroup hinting female mediated admixture instead of a male mediated admixture typical of most of the Indo-European castes of India.

## Material and Methods

### Sampling

Blood samples were collected from 213 individuals belonging to Nair, Thiyya, Bunt, Ezhava and Hoysala populations inhabited in Konkan and Malabar region of Karnataka and Kerala states in India (Figure 1a). All the subjects included in this study were healthy and unrelated. Informed written consent was obtained from each participant. The project was carried out in accordance with the guidelines approved by the Institutional Ethical Committees of Centre for Cellular and Molecular Biology, Hyderabad, India.

### Genotyping and Quality Control

DNA was isolated from the blood using standard phenol and chloroform method. We genotyped 76 samples on Affymetrix Axiome GW Human Origin Array for 633,994 SNPs as per the manufacturer’s specifications. Whole mitochondrial genome of all samples was also PCR amplified using a set of 24 markers and sequencing was done using Sanger sequencing method in ABI 3730 Automated DNA analyzer (Applied Biosystems, Foster city, USA). All mtDNA sequences were assembled with the revised Cambridge reference sequence (rCRS) (Andrews, et al. 1999) using AutoAssembler. Variations observed were used to assign the haplogroup using Phylotree build 17 (van Oven and Kayser 2009) and Haplogrep2 (Weissensteiner, et al. 2016).

For autosomal analysis, we merged our dataset with published DNA dataset (Reich, et al. 2009; Moorjani, et al. 2013; Mallick, et al. 2016; Nakatsuka, et al. 2017) of modern individuals after filtering for missingness using Plink 1.9 (Purcell, et al. 2007), and included only autosomal markers on 22 chromosomes having genotyping call rate > 99% and minor allele frequency > 1%. We further pruned dataset by removing individuals with first-degree and second-degree relatedness utilizing KING-robust (Manichaikul, et al. 2010) feature implemented in Plink2 (Chang, et al. 2015). After all the filtering, final merged dataset comprised of 968 contemporary individuals covering 422810 SNPs.

In order to minimize the effect of background LD in PCA and ADMIXTURE like analysis, we further thinned the markers by removing SNPs in strong LD (r2 > 0.4, window of 200 SNPs, sliding window of 25 SNPs at a time) using Plink 1.9 (Purcell, et al. 2007). For all the analyses with ancient DNA, we merged our samples with west Eurasian aDNA published datasets (Meyer, et al. 2012; Raghavan, et al. 2014; Allentoft, et al. 2015; Haak, et al. 2015; Mathieson, et al. 2015; Broushaki, et al. 2016; Lazaridis, et al. 2016; Yang, et al. 2017; Damgaard, et al. 2018; de Barros Damgaard, et al. 2018; Narasimhan, et al. 2019) of 1245 individuals which are relevant as reference for our population. In this aDNA merged dataset, we applied the missingness criteria of geno > 0.7, to include only those individuals covered for at least 70% of sites resulting into 1026 individuals covered at 441609 sites.

### Genome-wide SNP Data Analyses

We first performed Principal Component Analysis (PCA) on merged dataset of modern Eurasian using the *smartpca* package implemented in EIGENSOFT 7.2.1 (Patterson, et al. 2006) with default settings. We plotted first two components to infer genetic variability. We ran the model-based clustering algorithm ADMIXTURE (Alexander, et al. 2009) to infer ancestral genomic components in all five groups of South West coastal population inferred by PCA analysis. Cross validation was run 25 times for 12 ancestral clusters (K=3 to K=14). Lowest CV error parameter was obtained at K = 3. Therefore, we are showing the result for K value of 3. We constructed a maximum likelihood (ML) tree for our merged dataset of modern west Eurasian and south Asian populations using TreeMix v.1.12 (Pickrell and Pritchard 2012) with LD blocks of 500 SNPs grouped together and Onge as an outgroup.

We used ADMIXTOOLS 2 package in R to calculate Admixture F3 statistics and D-statistics and also to perform admixture modelling using qpAdm and qpGraph implementation. For the purpose of F3 statistics calculation and admixture graph, we used precalculated F2 statistics setting parameter maxmiss = 1, while for calculation of D-statistics and admixture modelling, we used genotype file directly and limiting the number of populations each time.

We used F3 statistics in the form of F3 (X, Palliyar; South West coast population), where X is any modern West Eurasian or South Asian population and Palliyar was used as a proxy for ASI ancestry. We calculated D-statistics (Patterson et al. 2012) in the form of Dstat (Nair/Bunt/Thiyya/Ezhava/Hoysala, X, Y; Yoruba) to infer the extent of gene flow into five population groups of South West coastal India. Where X is other South West coastal or Godavari region population and Y is North West India, North India, Pakistan and other west Eurasian populations.

We further utilized haplotype-based approach implemented in ChromoPainter (Lawson, et al. 2012) and fineSTRUCTURE (Lawson, et al. 2012) to derive co-ancestry matrix and fine scale population clustering, respectively. We first phased our data with SHAPEIT4 (Delaneau, et al. 2019) using default parameters. Chromosome painting was performed using ChromoPainter (Lawson, et al. 2012), first by performing 10 Expectation-Maximization (EM) iteration with 5 randomly selected chromosomes with a subset individuals to infer global mutation rate (μ) and switch rate parameters (Ne). Then, we ran the main algorithm with 22 chromosomes and including all the individuals to get coancestry matrix. This coancestry matrix was used by fineSTRUCTURE (Lawson, et al. 2012) to derive clustering using a probability model by applying Markov chain Monte Carlo (MCMC) procedure and then inferring hierarchical tree by merging all clusters with least change in posterior probability. For the run, we used 500,000 burn-in iterations and 1,000,000 subsequent iterations and stored the results from every 10,000^th^ iteration.

To visualize geographical population structure and diversity across India and compare it with South West coastal India, we ran Estimation of Effective Migration surface (EEMS) (Petkova, et al. 2016). For this, we first applied different number of demes ranging from 150 to 250 and also tuned proposal variances such as those were accepted 10-40% of times. For final run we chose 200 demes, with 10 million MCMC, 2 million burn-in, and 10,000 thinning iterations for the determination of posterior distribution of effective migration and effective diversity rates.

In order to gain further insight into temporal dynamics of effective migration and diversity rates, we used the tool MAPS (Al-Asadi, et al. 2019), which uses IBD sharing matrix and different length segments of IBD to track the change in effective migration and diversity rates with time. We used two different lengths intervals viz. 1-5 cm and 5-10 cm which corresponds to 90 generations and 22 generations ago, respectively. We used 200 demes with 5 million MCMC, 1 million burn-in and 10,000 thinning iterations for the determination of posterior distribution of effective migration and effective diversity rates. Here, again we tuned proposal variances for the acceptance proportions of 10-40% times.

We estimated the effective population size using Relate (Speidel, et al. 2019) package in R, which estimates genome wide genealogy applying Hidden Markov Model approach. First, assuming constant population size, it infers branch length estimates and then using Maximum Likelihood approach infers coalescence rate between pair of haplotypes. Inverse of the average value of coalescence rate gives the population size estimate. Repeated steps with MCMC convergence give us the joint estimates of branch lengths and effective population size. Here, we used all five study populations (Nair, Bunt, Thiyya, Ezhava and Hoysala) jointly as South West coast population along with two 1000 genome project populations (CEU and YRI), Iranian as Middle East group, Gangetic plain populations and populations from North West India and jointly inferred their effective population size and separation history. We also inferred signal of positive selection in above population groups by making use of genome-wide genealogy estimates by Relate (Speidel, et al. 2019).

In the analysis of merged dataset with ancient DNA references, we computed D-statistics in the form F4 (pop1, pop2, Steppe/Yamnaya, Yoruba) and F4 (pop1, pop2, Iran_N, Yoruba) to compare relative affinity of various modern Indian populations for Steppe vs Iranian ancestry in comparison to our study groups. Here, pop1 is various modern Indian cline groups and pop2 is our study group (Nair/Bunt/Thiyya/Ezhava/Hoysala) of the South West coastal India.

We used *qpadm* function of R package ADMIXTOOLS 2 to estimate proportions of ancient ancestral components in a test population (Nair/Bunt/Thiyya/Ezhava/Hoysala) derived from a set of N source population groups having shared drift with a set of reference populations. We performed modelling of admixture using Pre-bronze age and bronze age sources of Iranian ancestry viz Namazga_CA and Indus_Periphery group, respectively. We used Ethiopia_4500BP_published.SG, Anatolia_N, Dai.DG, Russia_EHG, WEHG, Jordan_PPNB, Ganj_Dareh_N, Israel_Natufian as references. We also obtained fitted admixture graph topology with *qpGraph* function In ADMIXTOOLS 2 package for five study groups along with Reddy population from Godavari basin and also Gujjar population from North West India and made a comparison with Gangetic plain population, Tiwari Brahmins.

## Supporting information

Supplimental Sheet

## Tools used

Admixtools 2: A new, lightning-fast implementation of ADMIXTOOLS.

https://github.com/uqrmaie1/admixtools **(manuscript describing the tool under preparation)**

## Availability of data and material

Data has been uploaded in the CCMB server http://tdb.ccmb.res.in/bic/database_pagelink.php?page=wcdata and would be made available for the researchers upon request to the corresponding authors.

## Acknowledgements

We thank all the study participants, who volunteered in this study. LK acknowledge the Council of Scientific and Industrial Research (CSIR) for the Fellowship. K.T was supported by the JC Bose Fellowship (JCB/2019/000027), SERB, Department of Science and Technology and Council of Scientific and Industrial Research, Ministry of Science and Technology, Government of India.

## Ethics declarations

### Conflict of interest

None.

### Ethics approval

The study was approved by Institute Ethical Committee of CSIR-CCMB, Hyderabad, India.

### Informed consent

Informed written consent was obtained from all the participants involved in the study.

## Notes

http://tdb.ccmb.res.in/bic/database_pagelink.php?page=wcdata

### Competing Interest Statement

The authors have declared no competing interest.

