## Supplementary material for "Genetic affinities and adaptation of the South West coast populations of India": Supplimental text

### Brief Introduction and history of south west coastal populations

#### Nairs

The Nayars were swordsmen, the military class of west coast of India. Such a description is found in **Pliny the Elder's records (Natural History VI Chapter 2) (Holland and Club 1848)**. Socially, the Nayars ranked after the Nambuthiris in Malabar. And occupied same position in the native states of Cochin and Travancore. **F. Fawcett in his book 'Nayars of Malabar' (1915) (Fawcett 1915)** emphasises matriarchy followed in this population and states – "The chief immediate interest attached to them lies in the fact of their being the best, that is the fullest, the most complete existing example of matriarchy, or, to be more strictly accurate, of inheritance through females. This system, obtaining at one time amongst the Celts and other races of Europe, was probably universal in the sense that it existed at some period in the life history of mankind, and is now to be found here and there in the world". He also goes on to speculate the origin of Nayars among the Aryan people as their appearance, customs, habitations and mode of life are very different from the other Dravidians of the Madras Presidency. However, he concludes "It is not yet time to say whether they are or are not (Dravidians)".

**K. Ananthakrishna Iyer, in his book 'Cochin Tribes and Castes' (1912) (Iyer 1969)**, describes them as the last of the honoured castes under the name of the pure "Sudras of Malayala". Nayars were mostly nobles who did not engage themselves in handicraft, commerce or any other occupation carried out by the Tiyans or Ezhavas. They instead carried arms and believed to have descended from a noble lineage.

The word Nayar is derived from Nayaka which means leader of the people. K. A. Iyer also notes that some people consider it to be more correct to derive Nayars from Nagas (Snakes or snake worshipping Nagas or Scythians). **Kerala Mahatmyan, a mythological and historical document in Sanskrit** assumes Nayars to be the descendants of the union of the junior members of the Nambuthiri families with the Devagandharva and Rakshasa womenbrought in by Parasurama from extra-terrestrial regions. Another ancient record **Keralotpatti** regards them as the descendants of the Sudras who accompanied the Brahman immigrants from outside Kerala. There are assumptions about their arrival from Tibet and Nepal. In the book **'The Sun and the Serpent' (1905) (Oldham 1905), C. F. Oldham** tries to draw a link between the serpent worshippers of the north and south specifically between the Nairs and the Newars of Nepal. Not just the appearance and tradition, Dravidian languages too retain a more intimate connection with the Scythian or Turanean tongues. He goes on to conclude that the Nayars, the original serpent worshipping population belonged to the Naga colonists of the North when the Southern confines of Central Asia was dominated by Dravidian (Asuras) kingdoms. However, there is a general consensus of opinion that Nayars are early immigrants from the Tamil country.

The Nayars are divided into clans, mostly exogamous with some endogamous groups. F. Fawcett also notes that the Nayar women of North Malabar were prohibited from intermingling with the men of South Malabar whom they considered inferior. This restriction was followed by the South Malabar clans too. The Nayars followed an interesting tradition which describes the social structure of the Malabar in the earlier times. The soiled cloth of the Nayar woman during her menstruation, as a rule, was washed by the washerwoman of the Tiyan caste and one of her own clean clothes were given to her instead. This was considered a purification process known as 'mattu' (change). This word is still in use in neighbouring Kannada and Tulu languages.

Nayars/Nairs at present constitute to 15-20% of the total population in Kerala (Various Surveys) with noticeable representation in the fields of Art, Science and Entrepreneurship.

#### Thiyya

Thiyya/Tiyans is a toddy-drawing caste from of Malabar, Cochin and Travancore. Until the 19th century this middle-class community was involved in any kind of work like cultivating the land, follow trades and professions, take service as domestics and so on. "Anything but soldiering, for which they have an utter abhorrence", Edgar Thurston states in his anthropological book series namely **'Castes and Tribes of Southern India' Volume VII published in 19' (Thurston and Rangachari 2018)**. During the British rule, there are official records stating European admixture which could be attributed to any Caucasian haplogroups found in the study.

It is believed that Tiyans and Izhuvans (another toddy drawing caste) migrated from Ceylon. **The South Canara Manual (Sturrock 1894)** states that "it is well known that both before and after the Christian era there were invasions and occupation on the northern part of Ceylon by the races then inhabiting Southern India, and Malabar tradition tells us that some of these Dravidians migrated again from Iram or Ceylon northwards to Travancore and other parts of the west coast of India, bringing with them the cocoanut or the southern tree (Tengina Mara) and being known as Tivars (Islanders) or Iravars, which names have been altered to Tiyars and Ilavars".

Socially, Tiyars who were considered to be subordinates of Nayars and Nambutiris, wouldn't eat rice cooked by an Izhuvan- pointing the inferiority of Izhuvans. Marriage is strictly prohibited between persons from the same illam (father's family). Each illam is exogamous. The main feature of the Tiyan religion is that it is largely connected to Shakti worship. Arrack is prominently used in ceremonies instead of milk or honey. There are a lot of similarities between the cultural and social practices of Tiyans, Ilavas (Izhuvans), Billavas and Poojaris. Sri Narayan Guru is revered by the people of all these communities. It is to be noted that, these communities are matrilineal. However, **A Shreedhar Menon in his 'A survey of Kerala History' (Menon 2006)**, opines that Marumakkattayam, matrilineal system was introduced into the Malabar region after the Chola Chera war during the eleventh century A.D. In his thesis titled **'The History of the Awakening of the Ilava community in Kerala' (Jacob 1982), George Jacob** groups Tiyans as subset of the larger modern day Ilava community, limited to the North Malabar region**. A. Aiyyappan in his book 'Iravas and culture change' (Aiyappan 1944)** describes 'Tiya' as a synonym for Iravas/Ilav/Izhuvans in North Malabar, 'Tandan' in Walluvanad taluk of South Malabar and 'Chovan' in southern divisions in Cochin. He also states that since Tiyas are educationally and economically more advanced, at every census, Iravas and Tandans identify themselves as Tiyas.

The legends, however, have differentiated Tiyas and Iravas distinctly. Iravas originally came from Ceylon and settled in the Malabar region. An ancient king of Malabar insulted the artisan section of the population, who left his kingdom and settled in Ceylon. On their return they were escorted by the 'protectors' who the Iravas claim to be their ancestors. One of the well-known legends classifies Tiyas and Tandans as descendants of the returned Iravas from Ceylon and the local Brahman woman. While the Irava Pannikars descended from a Chruma woman.

#### Bunts

Bunt is a warrior class community in the coastal districts of Karnataka speaking Tulu as well as Kundagannada as their mother tongue and were traditionally an agrarian caste engaged in rice cultivation. The Bunts follow a matrilineal system of inheritance called Aliyasantana. They have 93 clan names or surnames and are divided into 53 matrilineal septs called Bari. Members of the same bari did not intermarry. According to description by Historian Edgar Thurston in his book "Caste and tribes of south India" (1909) - "Men and women of the Bunt community belong to a beautiful race of Asia. Men have a broad forehead and a parrot nose. Mostly they are of fair complexion. Even today they are of independent nature, short tempered, self-respecting and have a muscular body, which tells about the history of belonging to warrior families" (Thurston and Rangachari 2018).

Bunts relates their descent from the ancient Alupa dynasty (200 CE - 1500 CE) and according to Historian P. Gururaja Bhat the Alupa royal family were possibly belonging to the Bunt caste. According to Indian anthropologist Ayinapalli Aiyappan warrior clan of Kolathunadu (Kola Bari and the Kolathiri Raja) was a descendant of the Bunts. As mentioned by Norwegian anthropologist Harald Tambs-Lyche Jain kingdoms of Canara region during the Hoysala dynasty had warriors from Bunt clans and also Bunts were appointed as military officers under The Hoysala Ballal kings.

#### Ezhava

Ezhava is a population of Malabar coast of India constituting around 23% of the population of the area. They are social groups of agricultural labourers, small cultivators, toddy tappers along with involovement in weaving and also practicing Ayurveda. According to one theory based on numerous shared customs like childbirth, death, their matrilineal practices and martial history, they share common heritage with the Nairs.

Ezhava also worked as warrior’s clan under local rulers such as of Kadathanad and Kurumbranad of Kerala. A subgroup of the Ezhavas considered themselves to be warriors and became known as the Chekavars. The Vadakkan Pattukal ballads describe Chekavars as forming the militia of local chieftains and kings but the title was also given to experts of Kalari Payattu.

### Hoysala

Hoysala are one of the prominent Brahmin communities of Karnataka speaking Kannada, inhabiting the Southern Districts of Karnataka such as Shivamogga, Davanagere, Chitradurga, Chikmagalur, Hassan, Tumkur, Mysore, Mandya, Bangalore and Kolar. According to historians they have been natives of the Malnad region of Karnataka but inscriptions also point towards their contacts with Yadava from north India. During 10$^{th}$ century to 14$^{th}$ century, they ruled a large region of Karnataka, parts of Andhra Pradesh and Tamil Nadu as Hoysala dynasty. This dynasty was founded by King Nripa Kama ii, who was having an alliance with the western Ganga dynasty. Although proper records are lacking to link Hoysala to the Yadavas from north India.

### Supplementary Figures

#### Fig S1a-e. Admixture F3 statistics of south west coastal populations.

**A.** Admixture F3 statistics in the form F3(Nair; Palliyar, X), where X is any modern west Eurasians. **B.** Admixture F3 statistics in the form F3(Thiyya; Palliyar, X), where X is any modern west Eurasians. **C.** Admixture F3 statistics in the form F3(Bunt; Palliyar, X), where X is any modern west Eurasians. **D.** Admixture F3 statistics in the form F3(Ezhava; Palliyar, X), where X is any modern west Eurasians. **E.** Admixture F3 statistics in the form F3(Hoyslal; Palliyar, X), where X is any modern west Eurasians.

A

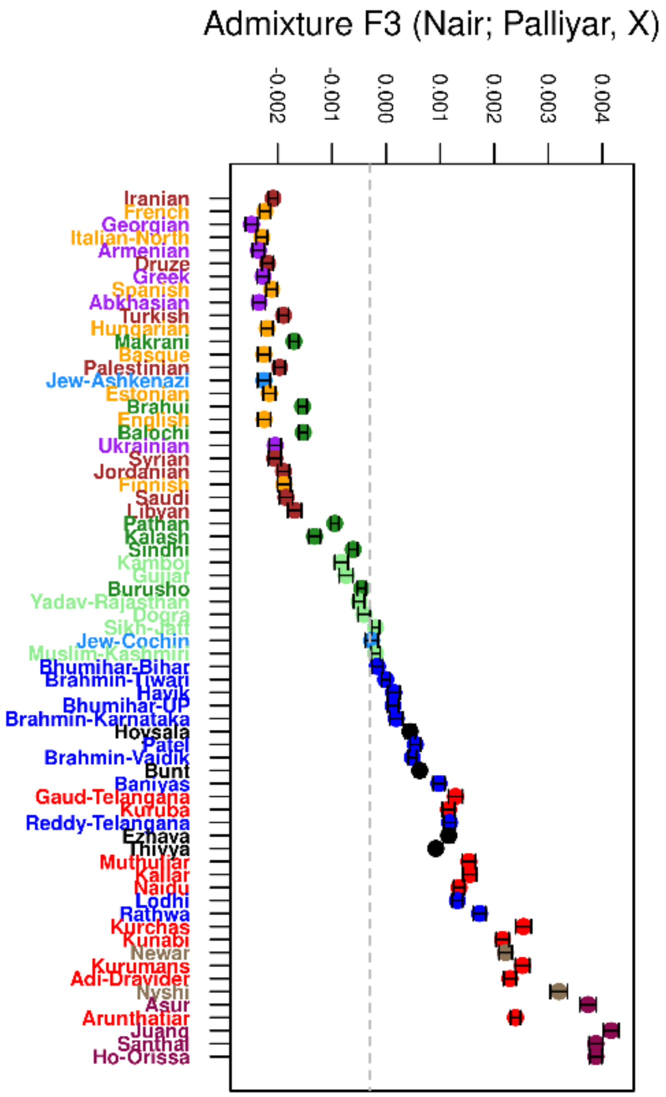

B

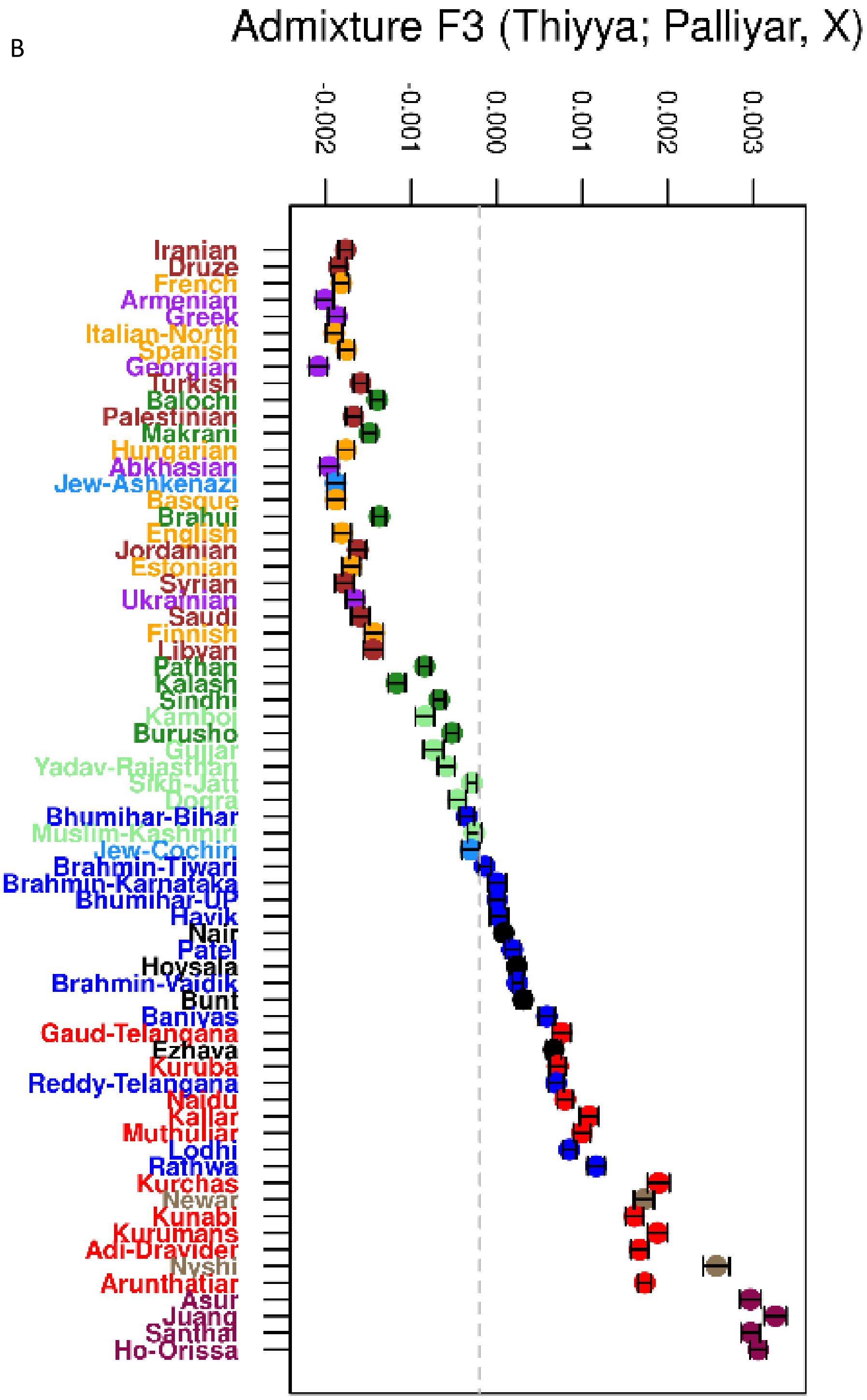
Admixture F3 (Thiyya; Palliyar, X)
-0.002
-0.001
0.000
0.001
0.002
0.003
Iranian
Druze
French
Armenian
Greek
Italian-North
Spanish
Georgian
Turkish
Balochi
Palestinian
Makrani
Hungarian
Abkhasian
Jew-Ashkenazi
Basque
Brahui
English
Jordanian
Estonian
Syrian
Ukrainian
Saudi
Finnish
Libyan
Pathan
Kalash
Sindhi
Kamboj
Burusho
Gujjar
Yadav-Rajasthan
Sikh-Jatt
Dogra
Bhumihar-Bihar
Muslim-Kashmiri
Jew-Cochin
Brahmin-Tiwari
Brahmin-Karnataka
Bhumihar-UP
Havik
Nair
Patel
Hoysala
Brahmin-Vaidik
Bunt
Baniyas
Gaud-Telangana
Ezhava
Kuruba
Reddy-Telangana
Naidu
Kallar
Muthuliar
Lodhi
Rathwa
Kurchas
Newar
Kunabi
Kurumans
Adi-Dravider
Nyshi
Arunthatiar
Asur
Juang
Santhal
Ho-Orissa

c

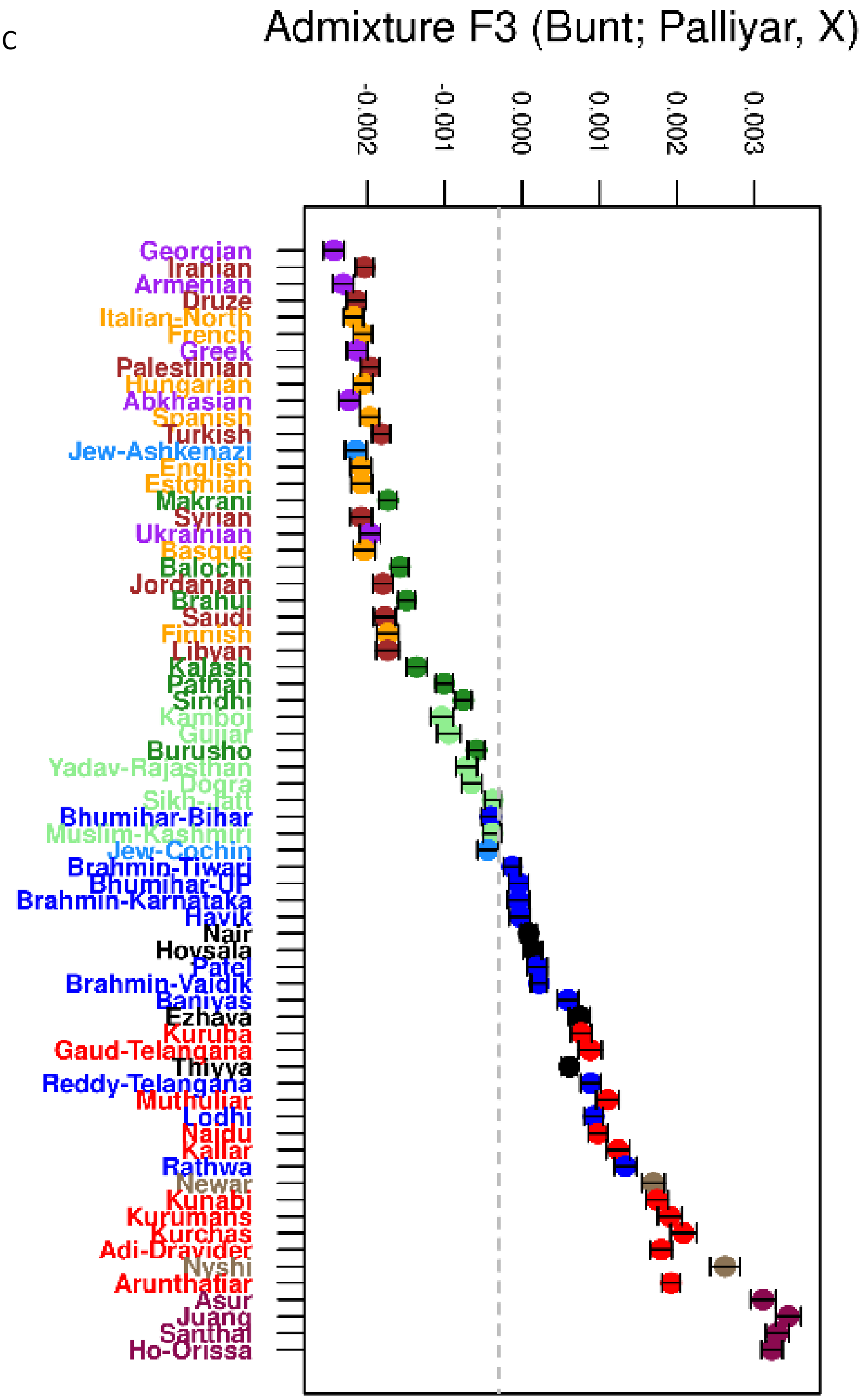
Admixture F3 (Bunt; Palliyar, X)
-0.002
-0.001
0.000
0.001
0.002
0.003
Georgian
Iranian
Armenian
Druze
Italian-North
French
Greek
Palestinian
Hungarian
Abkhasian
Spanish
Turkish
Jew-Ashkenazi
English
Estonian
Makrani
Syrian
Ukrainian
Basque
Balochi
Jordanian
Brahui
Saudi
Finnish
Libyan
Kalash
Pathan
Sindhi
Kamboj
Gujjar
Burusho
Yadav-Rajasthan
Dogra
Sikh-Jatt
Bhumihar-Bihar
Muslim-Kashmiri
Jew-Cochin
Brahmin-Tiwari
Bhumihar-UP
Brahmin-Karnataka
Havik
Nair
Hoysala
Patel
Brahmin-Vaidik
Baniyas
Ezhava
Kuruba
Gaud-Telangana
Thiyya
Reddy-Telangana
Muthuliar
Lodhi
Naidu
Kallar
Rathwa
Newar
Kunabi
Kurumans
Kurchas
Adi-Dravider
Nyshi
Arunthatiar
Asur
Juang
Santhal
Ho-Orissa

D

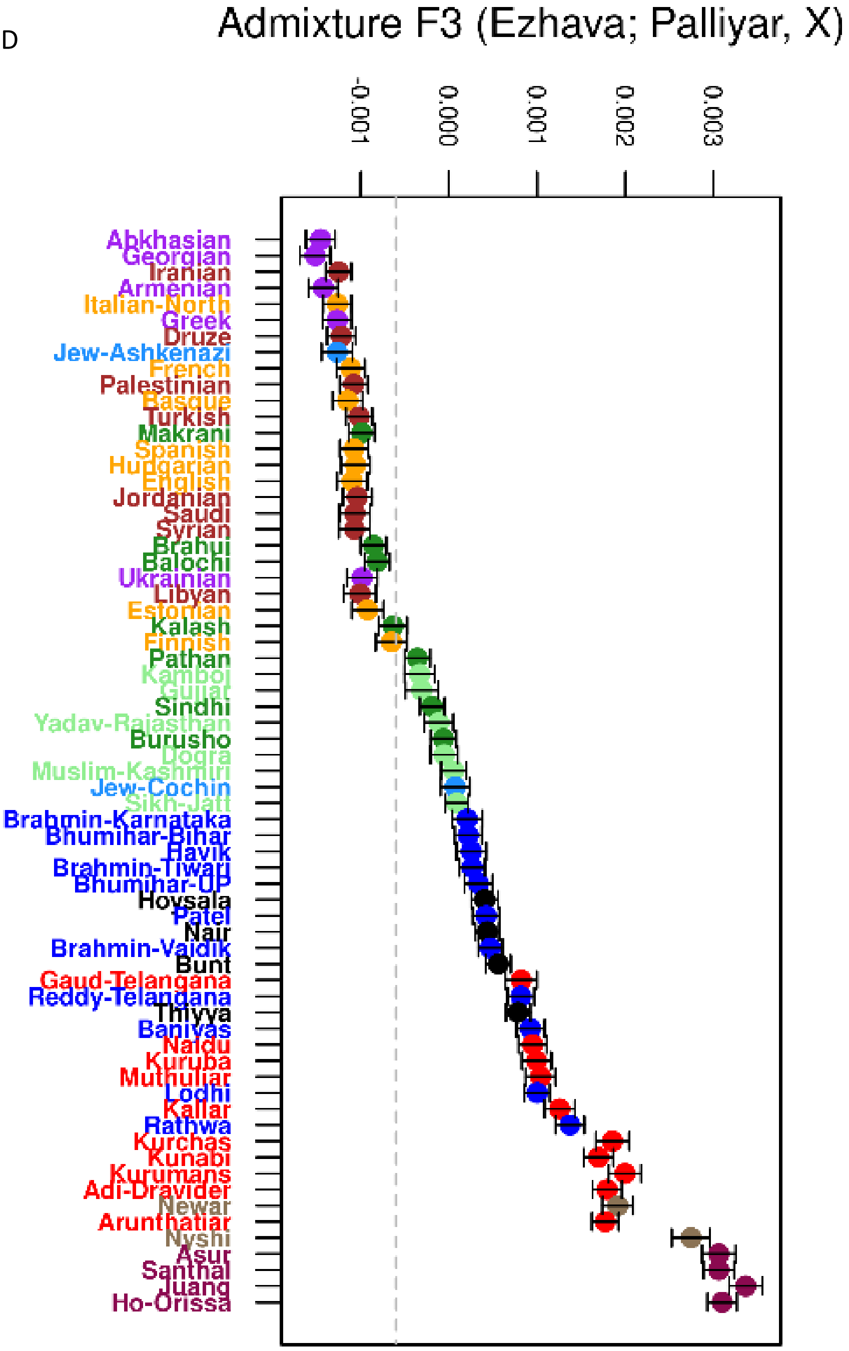
Admixture F3 (Ezhava; Palliyar, X)
-0.001
0.000
0.001
0.002
0.003
Abkhasian
Georgian
Iranian
Armenian
Italian-North
Greek
Druze
Jew-Ashkenazi
French
Palestinian
Basque
Turkish
Makrani
Spanish
Hungarian
English
Jordanian
Saudi
Syrian
Brahui
Balochi
Ukrainian
Libyan
Estonian
Kalash
Finnish
Pathan
Kamboj
Gujjar
Sindhi
Yadav-Rajasthan
Burusho
Dogra
Muslim-Kashmiri
Jew-Cochin
Sikh-Jatt
Brahmin-Karnataka
Bhumihar-Bihar
Havik
Brahmin-Tiwari
Bhumihar-UP
Hoysala
Patel
Nair
Brahmin-Vaidik
Bunt
Gaud-Telangana
Reddy-Telangana
Thiyya
Baniyas
Naidu
Kuruba
Muthuliar
Lodhi
Kallar
Rathwa
Kurchas
Kunabi
Kurumans
Adi-Dravider
Newar
Arunthatiar
Nyshi
Asur
Santhal
Juang
Ho-Orissa

E

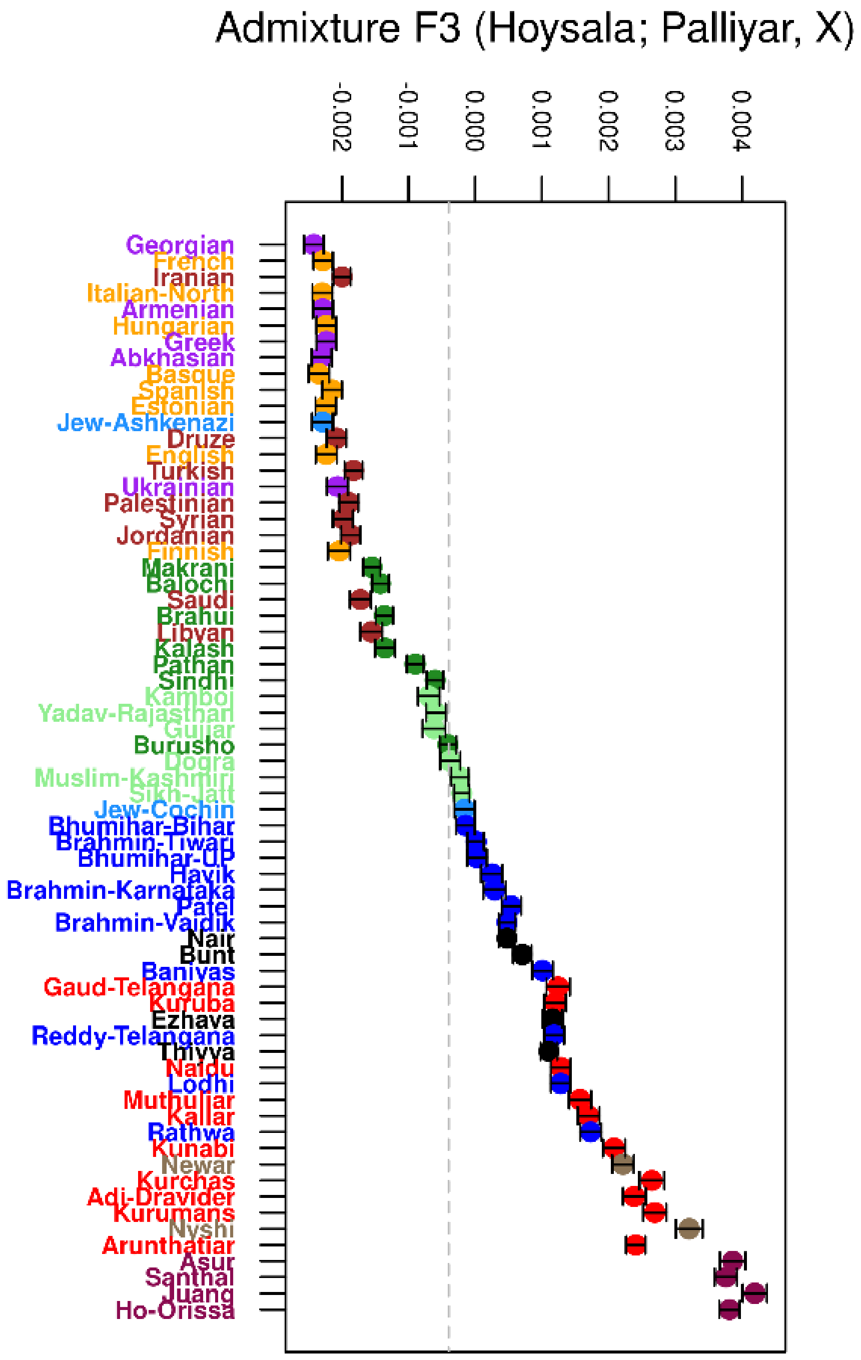
Admixture F3 (Hoysala; Palliyar, X)
-0.002
-0.001
0.000
0.001
0.002
0.003
0.004
Georgian
French
Iranian
Italian-North
Armenian
Hungarian
Greek
Abkhasian
Basque
Spanish
Estonian
Jew-Ashkenazi
Druze
English
Turkish
Ukrainian
Palestinian
Syrian
Jordanian
Finnish
Makrani
Balochi
Saudi
Brahui
Libyan
Kalash
Pathan
Sindhi
Kamboj
Yadav-Rajasthan
Gujjar
Burusho
Dogra
Muslim-Kashmiri
Sikh-Jatt
Jew-Cochin
Bhumihar-Bihar
Brahmin-Tiwari
Bhumihar-UP
Havik
Brahmin-Karnataka
Patel
Brahmin-Vaidik
Nair
Bunt
Baniyas
Gaud-Telangana
Kuruba
Ezhava
Reddy-Telangana
Thiyya
Naidu
Lodhi
Muthuliar
Kallar
Rathwa
Kunabi
Newar
Kurchas
Adi-Dravider
Kurumans
Nyshi
Arunthatiar
Asur
Santhal
Juang
Ho-Orissa

**Fig S2. *D* statistics of south west coastal population with ancient sources. A.** Steppe_EMBA vs. Iran_N scatterplot comparing D(X, Nair; Yamnaya, Yoruba) vs D(X, Nair; Iran_N, Yoruba), where X is different modern west Eurasian groups. **B.** Steppe_EMBA vs. Iran_N scatterplot comparing D(X, Thiyya; Yamnaya, Yoruba) vs D(X, Thiyya; Iran_N, Yoruba). **C.** Steppe_EMBA vs. Iran_N scatterplot comparing D(X, Bunt; Yamnaya, Yoruba) vs D(X, Bunt; Iran_N, Yoruba). **D.** Steppe_EMBA vs. Iran_N scatterplot comparing D(X, Ezhava; Yamnaya, Yoruba) vs D(X, Ezhava; Iran_N, Yoruba). **E.** Steppe_EMBA vs. Iran_N scatterplot comparing D(X, Hoysala; Yamnaya, Yoruba) vs D(X, Hoysala; Iran_N, Yoruba).

A

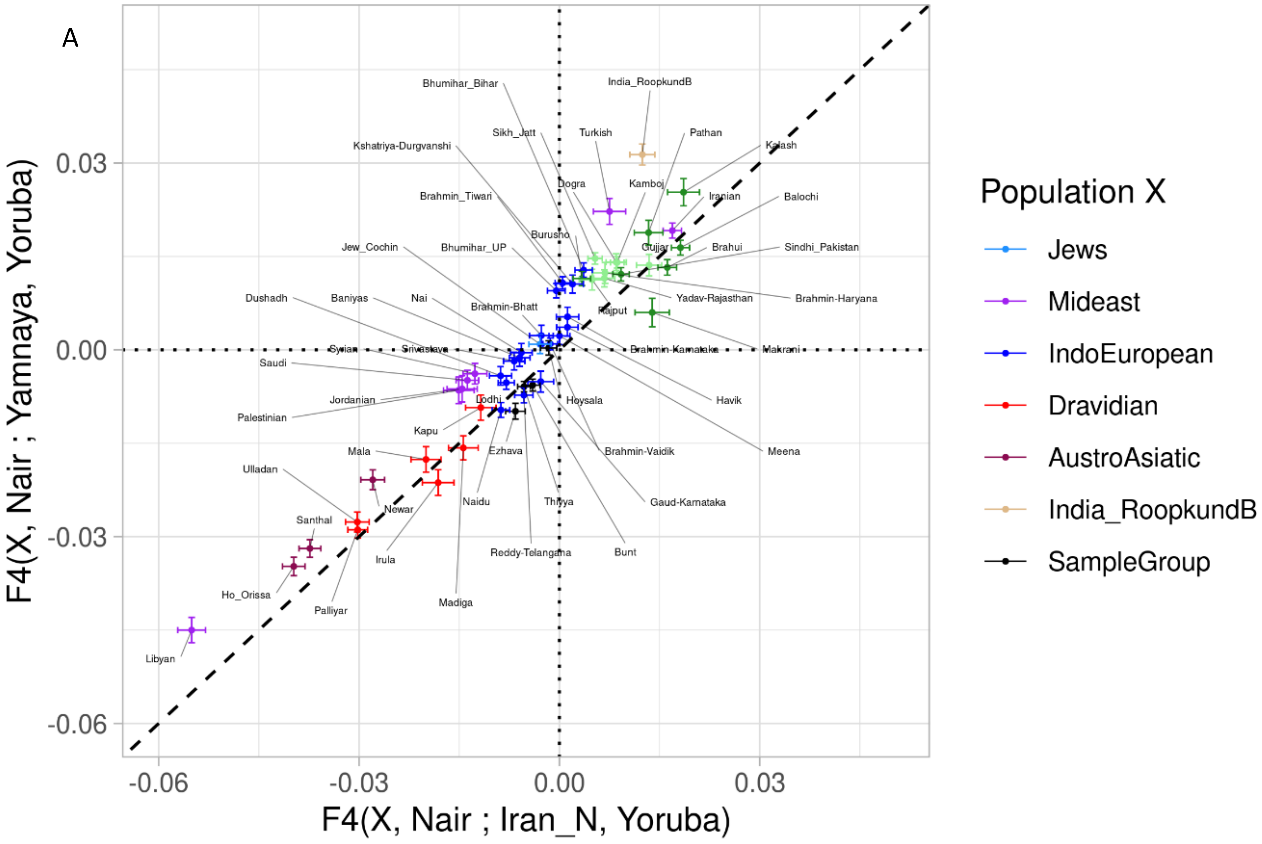

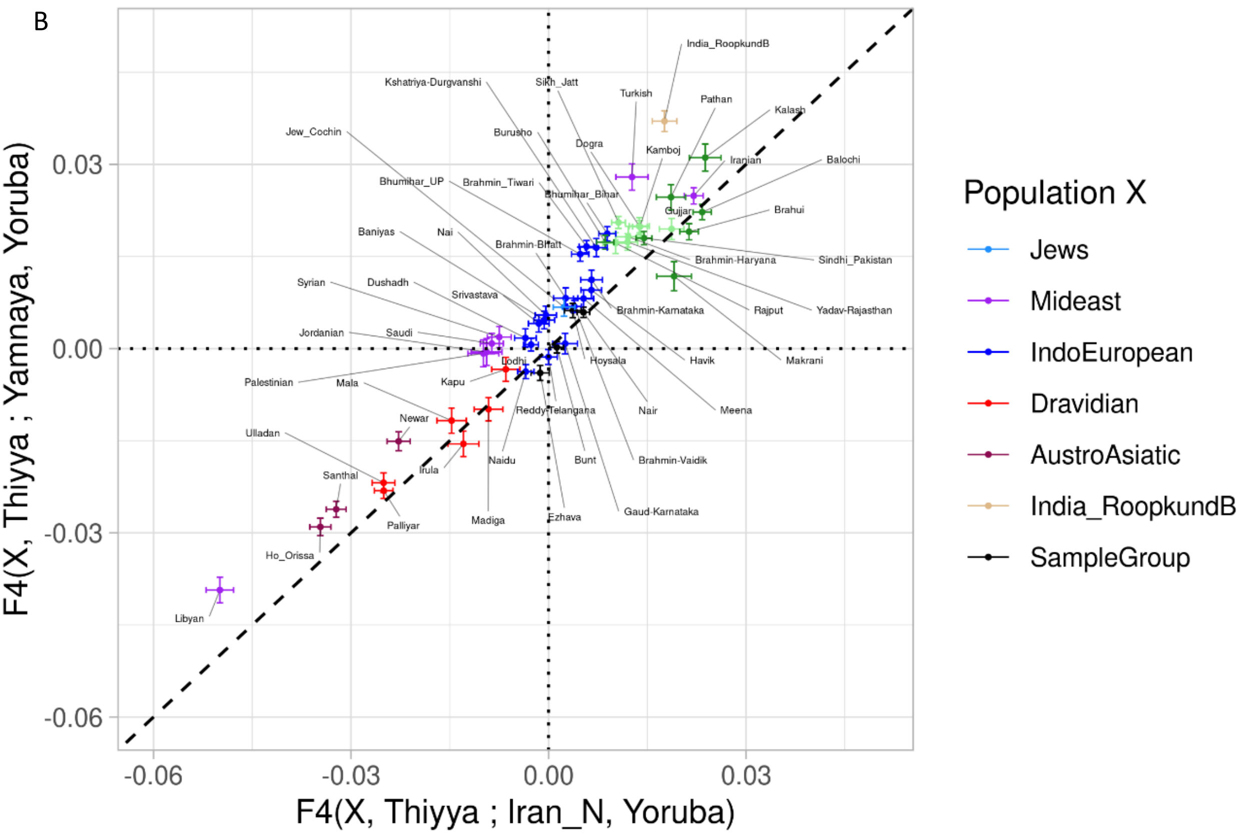
B
F4(X, Thiyya ; Yamnaya, Yoruba)
F4(X, Thiyya ; Iran_N, Yoruba)
0.03
0.00
-0.03
-0.06
Population X
Jews
Mideast
IndoEuropean
Dravidian
AustroAsiatic
India_RoopkundB
SampleGroup
India_RoopkundB
Kshatriya-Durgvanshi
Sikh_Jatt
Turkish
Pathan
Kalash
Jew_Cochin
Burusho
Dogra
Kamboj
Iranian
Balochi
Bhumihar_UP
Brahmin_Tiwari
Bhumihar_Bihar
Gujjar
Brahui
Baniyas
Nai
Brahmin-Bhatt
Brahmin-Haryana
Sindhi_Pakistan
Syrian
Dushadh
Srivastava
Brahmin-Karnataka
Rajput
Yadav-Rajasthan
Jordanian
Saudi
Lodhi
Hoysala
Havik
Makrani
Palestinian
Mala
Kapu
Reddy-Telangana
Nair
Meena
Newar
Ulladan
Naidu
Bunt
Brahmin-Vaidik
Irula
Santhal
Madiga
Ezhava
Gaud-Karnataka
Palliyar
Ho_Orissa
Libyan

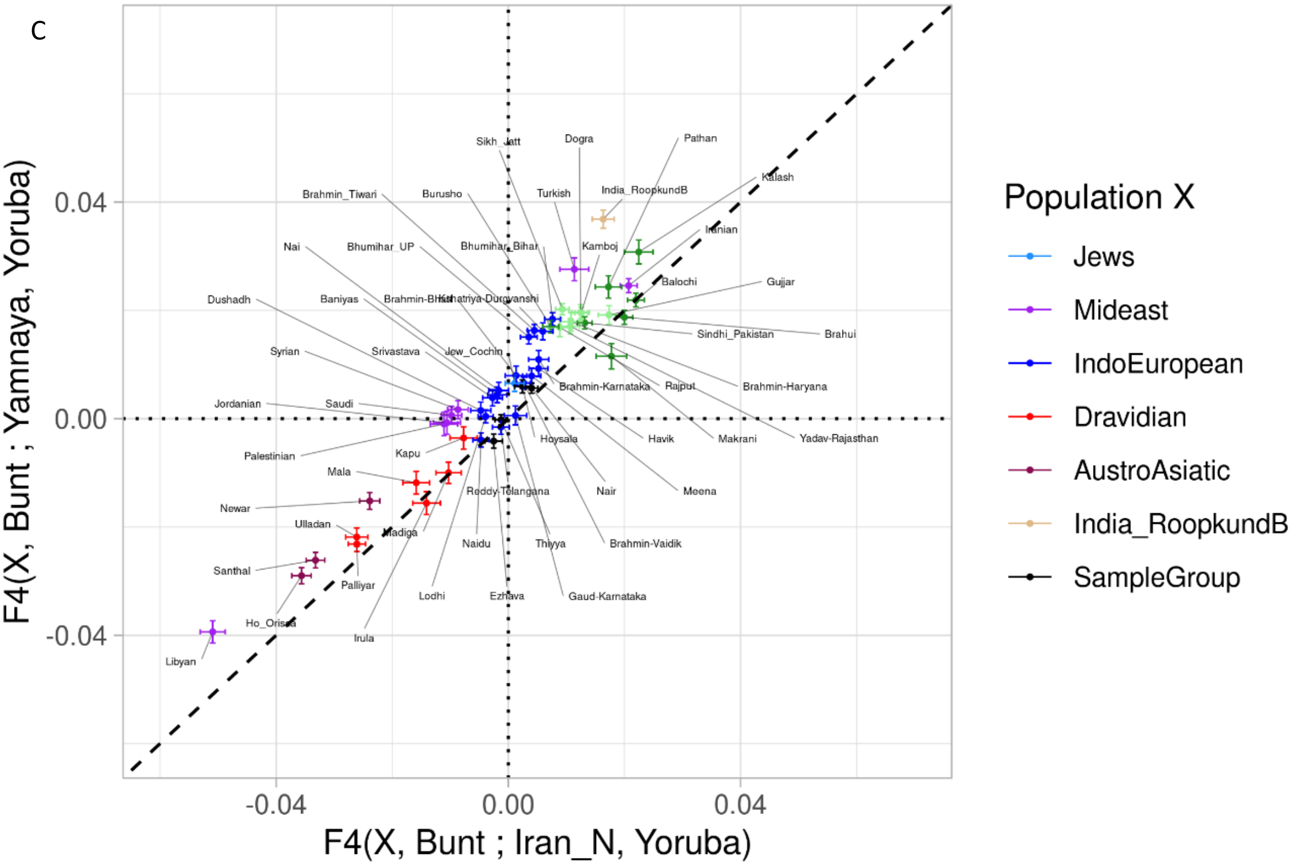
C
F4(X, Bunt ; Yamnaya, Yoruba)
F4(X, Bunt ; Iran_N, Yoruba)
0.04
0.00
-0.04
-0.04
0.00
0.04
Sikh_Jatt
Dogra
Pathan
Kalash
Brahmin_Tiwari
Burusho
Turkish
India_RoopkundB
Iranian
Nai
Bhumihar_UP
Bhumihar_Bihar
Kamboj
Balochi
Gujjar
Dushadh
Baniyas
Brahmin-Bihar
Kshatriya-Durgvanshi
Sindhi_Pakistan
Brahui
Syrian
Srivastava
Jew_Cochin
Brahmin-Karnataka
Rajput
Brahmin-Haryana
Jordanian
Saudi
Hoysala
Havik
Makrani
Yadav-Rajasthan
Palestinian
Kapu
Mala
Reddy-Telangana
Nair
Meena
Newar
Ulladan
Madiga
Naidu
Thiyya
Brahmin-Vaidik
Santhal
Palliyar
Lodhi
Ezhava
Gaud-Karnataka
Ho_Orissa
Irula
Libyan
Population X
Jews
Mideast
IndoEuropean
Dravidian
AustroAsiatic
India_RoopkundB
SampleGroup

D

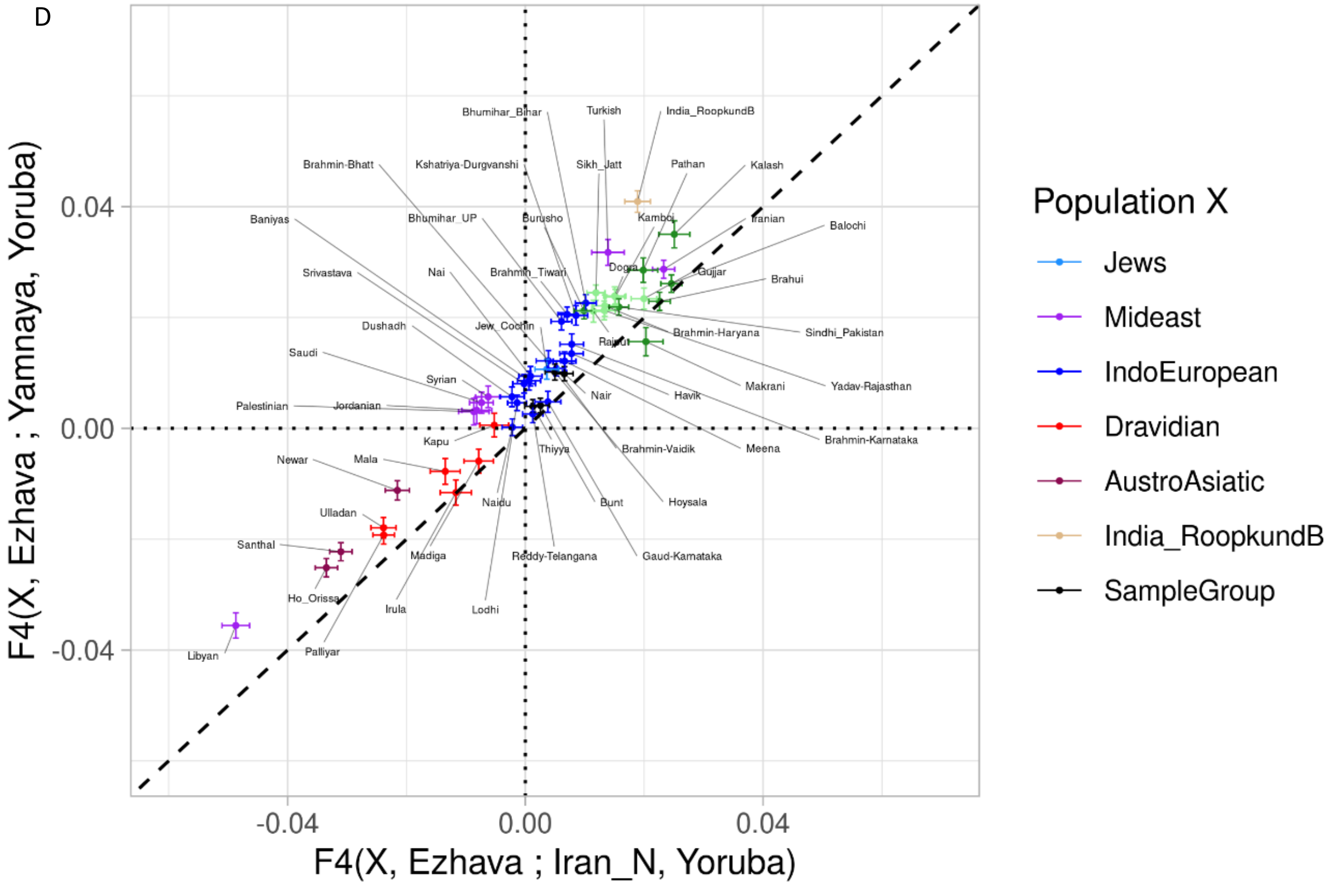
Bhumihar_Bihar
Turkish
India_RoopkundB
Brahmin-Bhatt
Kshatriya-Durgvanshi
Sikh_Jatt
Pathan
Kalash
Baniyas
Bhumihar_UP
Burusho
Kamboj
Iranian
Balochi
Srivastava
Nai
Brahmin_Tiwari
Dogra
Gujjar
Brahui
Dushadh
Jew_Cochin
Rajput
Brahmin-Haryana
Sindhi_Pakistan
Saudi
Syrian
Nair
Havik
Makrani
Yadav-Rajasthan
Palestinian
Jordanian
Kapu
Thiyya
Brahmin-Vaidik
Meena
Brahmin-Karnataka
Newar
Mala
Naidu
Bunt
Hoysala
Ulladan
Santhal
Madiga
Reddy-Telangana
Gaud-Karnataka
Ho_Orissa
Irula
Lodhi
Libyan
Palliyar
0.04
0.00
-0.04
-0.04
0.00
0.04
F4(X, Ezhava ; Yamnaya, Yoruba)
F4(X, Ezhava ; Iran_N, Yoruba)
Population X
Jews
Mideast
IndoEuropean
Dravidian
AustroAsiatic
India_RoopkundB
SampleGroup

E

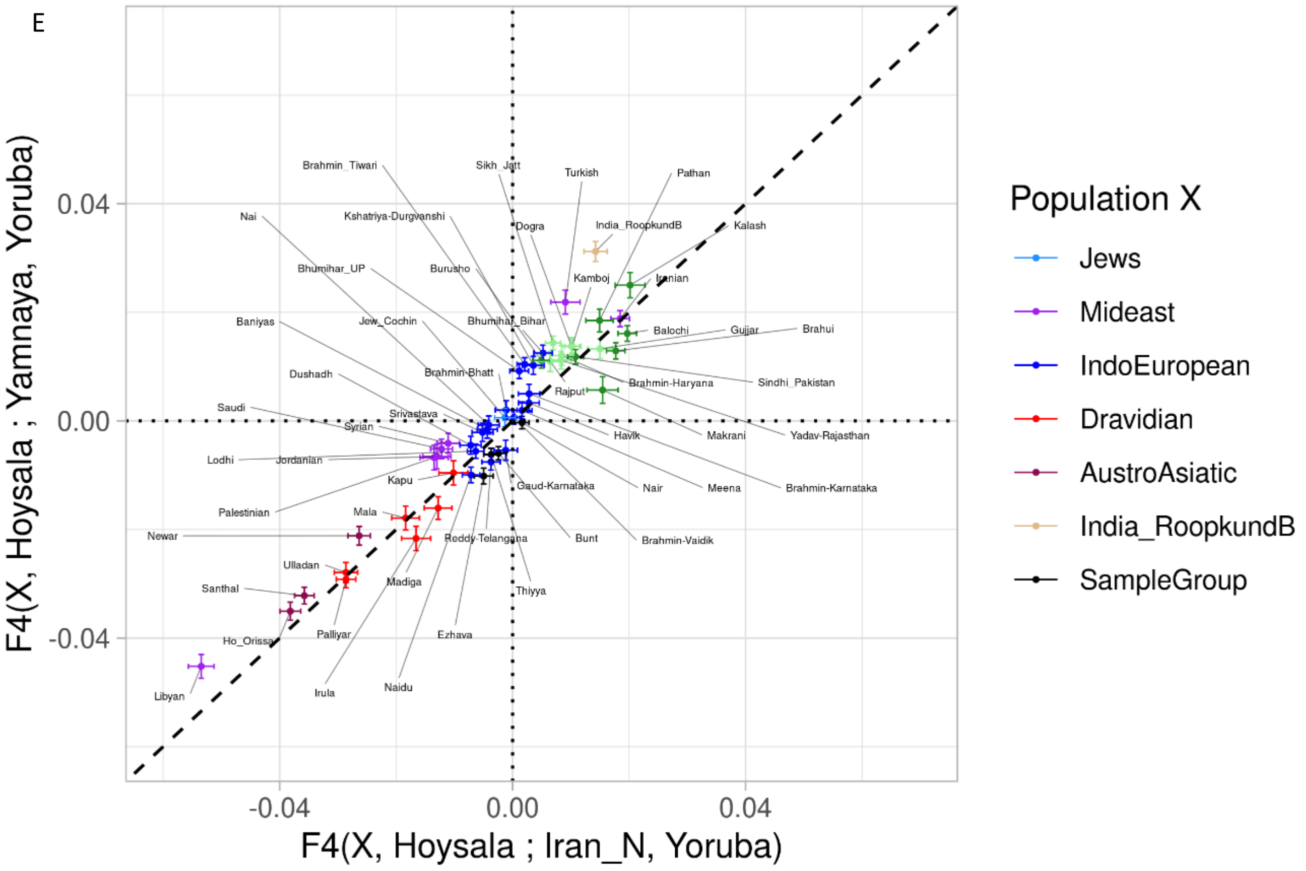
F4(X, Hoysala ; Yamnaya, Yoruba)
F4(X, Hoysala ; Iran_N, Yoruba)
0.04
0.00
-0.04
Population X
Jews
Mideast
IndoEuropean
Dravidian
AustroAsiatic
India_RoopkundB
SampleGroup
Brahmin_Tiwari
Sikh_Jatt
Turkish
Pathan
Nai
Kshatriya-Durgvanshi
Dogra
India_RoopkundB
Kalash
Bhumihar_UP
Burusho
Kamboj
Iranian
Baniyas
Jew_Cochin
Bhumihar_Bihar
Balochi
Gujjar
Brahui
Dushadh
Brahmin-Bhatt
Brahmin-Haryana
Sindhi_Pakistan
Rajput
Saudi
Srivastava
Syrian
Havik
Makrani
Yadav-Rajasthan
Lodhi
Jordanian
Kapu
Gaud-Karnataka
Nair
Meena
Brahmin-Karnataka
Palestinian
Mala
Newar
Reddy-Telangana
Bunt
Brahmin-Vaidik
Ulladan
Madiga
Thiyya
Santhal
Palliyar
Ezhava
Ho_Orissa
Libyan
Irula
Naidu

**Fig S3. Maximum Likelihood tree using TreeMix**: relationship of south west coastal groups with other south Asian and west Eurasians inferred by TreeMix.

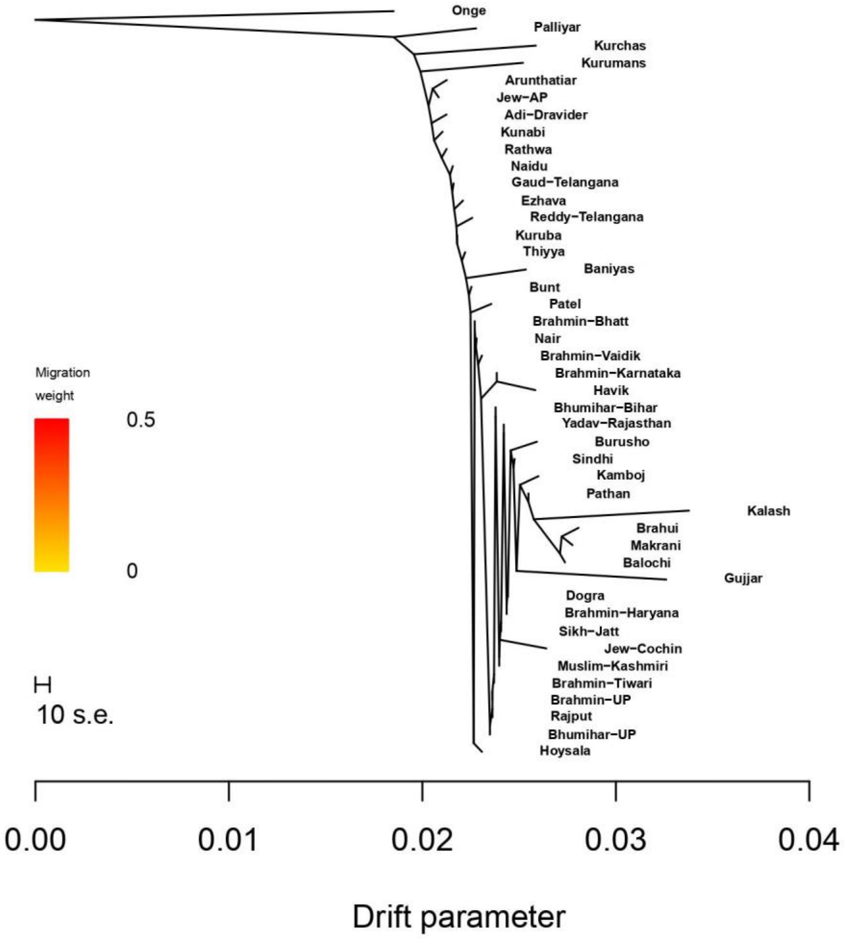

**Fig S4a.** Individual level phylogenetic tree for south west coastal groups along with other Indian and Pakistani groups based on haplotype sharing and constructed by FineStructure

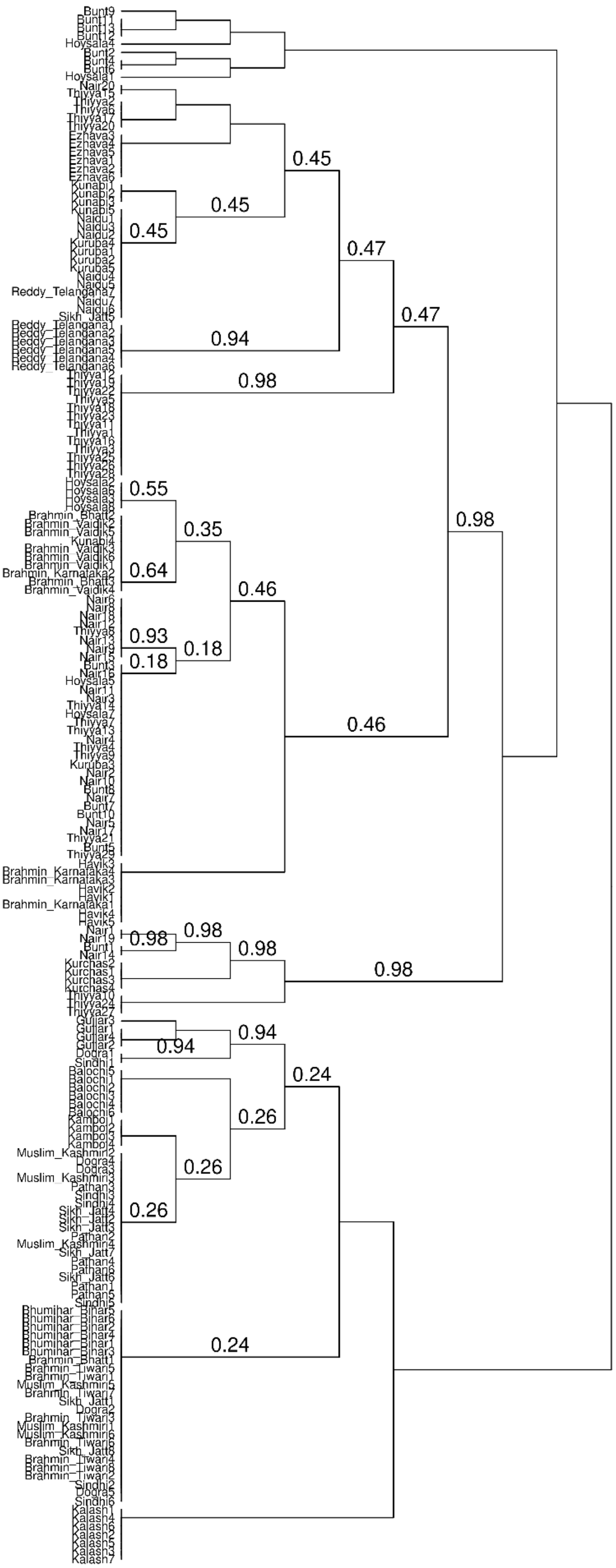

**Fig S4b.** Population level phylogenetic tree for south west coastal groups along with other Indian and Pakistani groups based on haplotype sharing and constructed by fineStructure

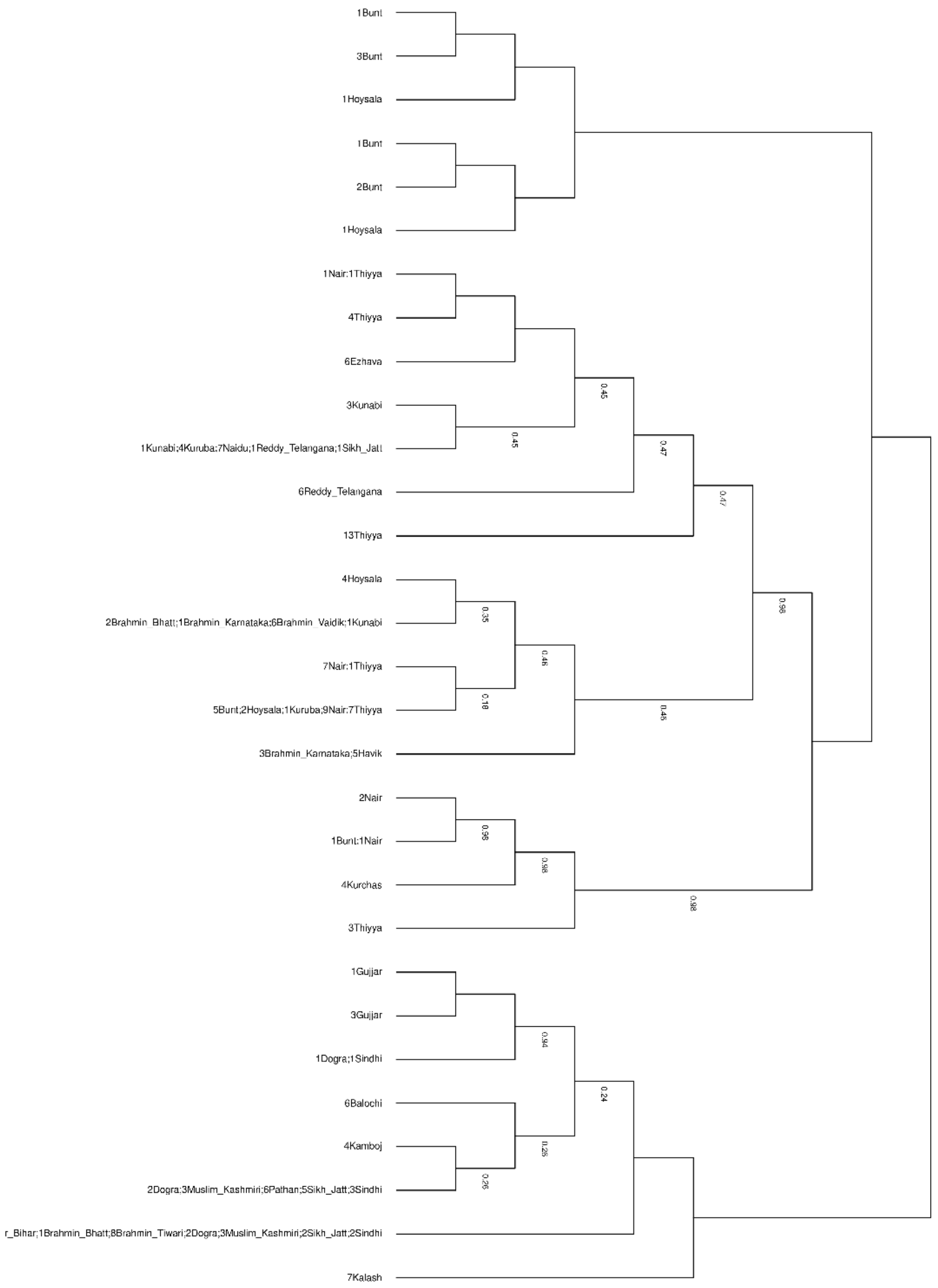

**Fig S5. Fitted admixture graph topology using qpGraph function in Admixtools 2.**

**A.** Fitted graph topology for Nair with ancient and some modern source groups showing additional Iranian component required from a distant ancestral source group related to BMAC. **B.** Fitted graph topology for Thiyya with ancient and some modern source groups. **C.** Fitted graph topology for Bunt with ancient and some modern source groups. **D.** Fitted graph topology for Ezhava with ancient and some modern source groups. **E.** Fitted graph topology for Gujjar with ancient and some modern source groups.

A

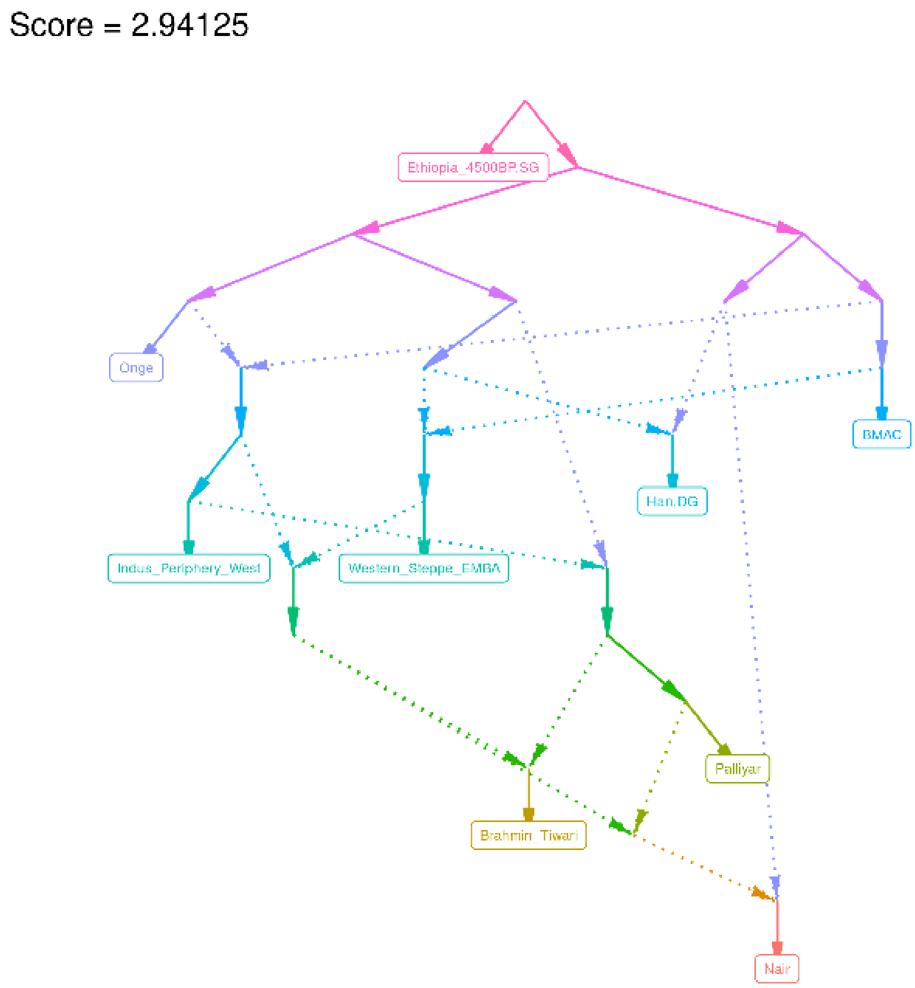

B

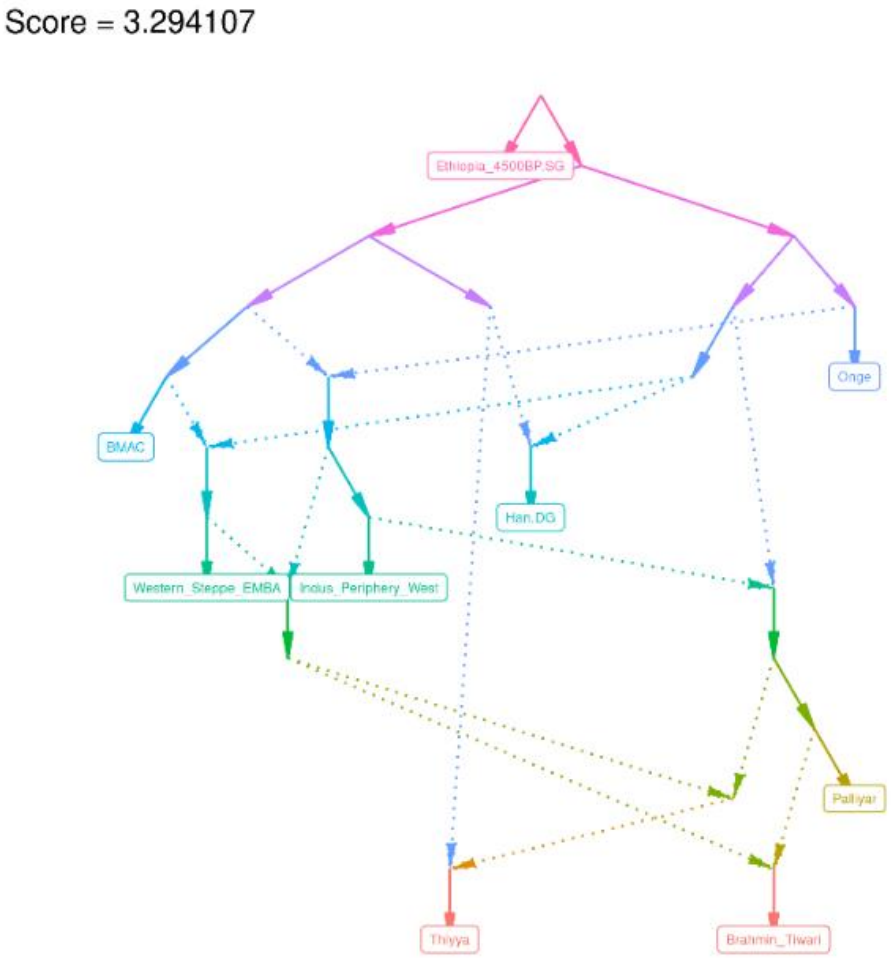
Score = 3.294107
Ethiopia_4500BP.SG
Onge
BMAC
Han.DG
Western_Steppe_EMBA
Indus_Periphery_West
Palliyar
Thiyya
Brahmin_Tiwari

C

Score = 2.919086

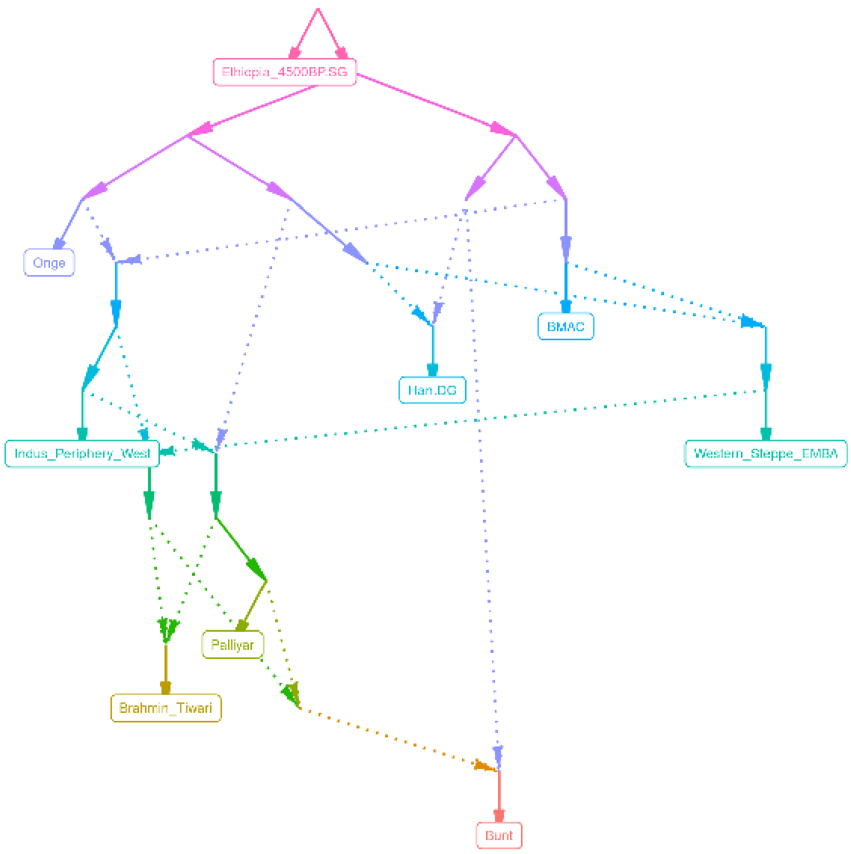

D Score = 2.817161

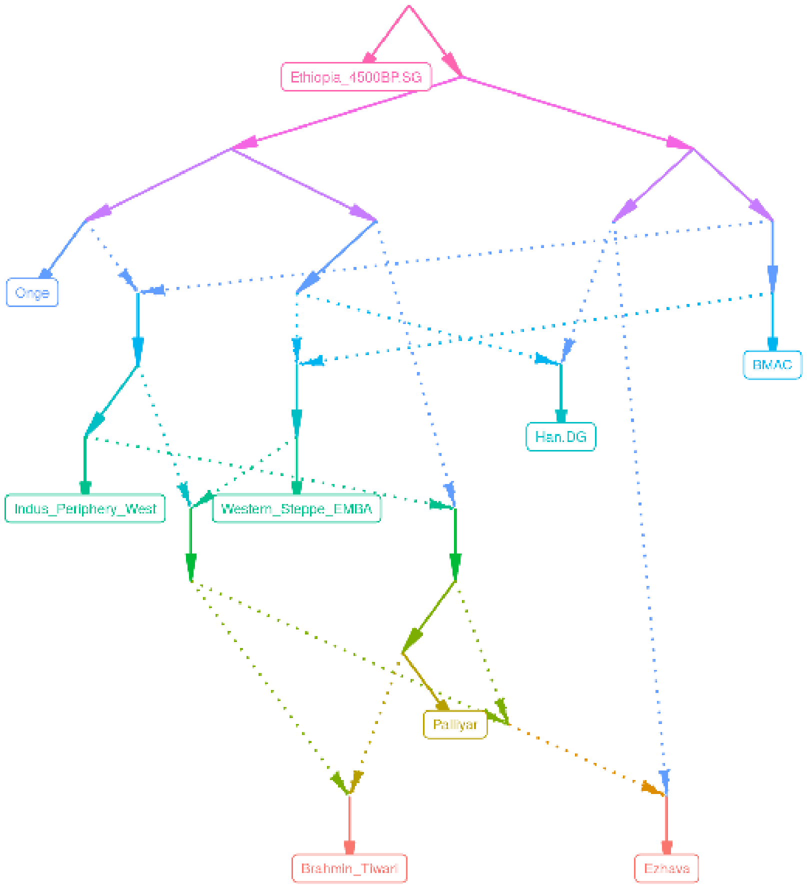

E Score = 2.307357

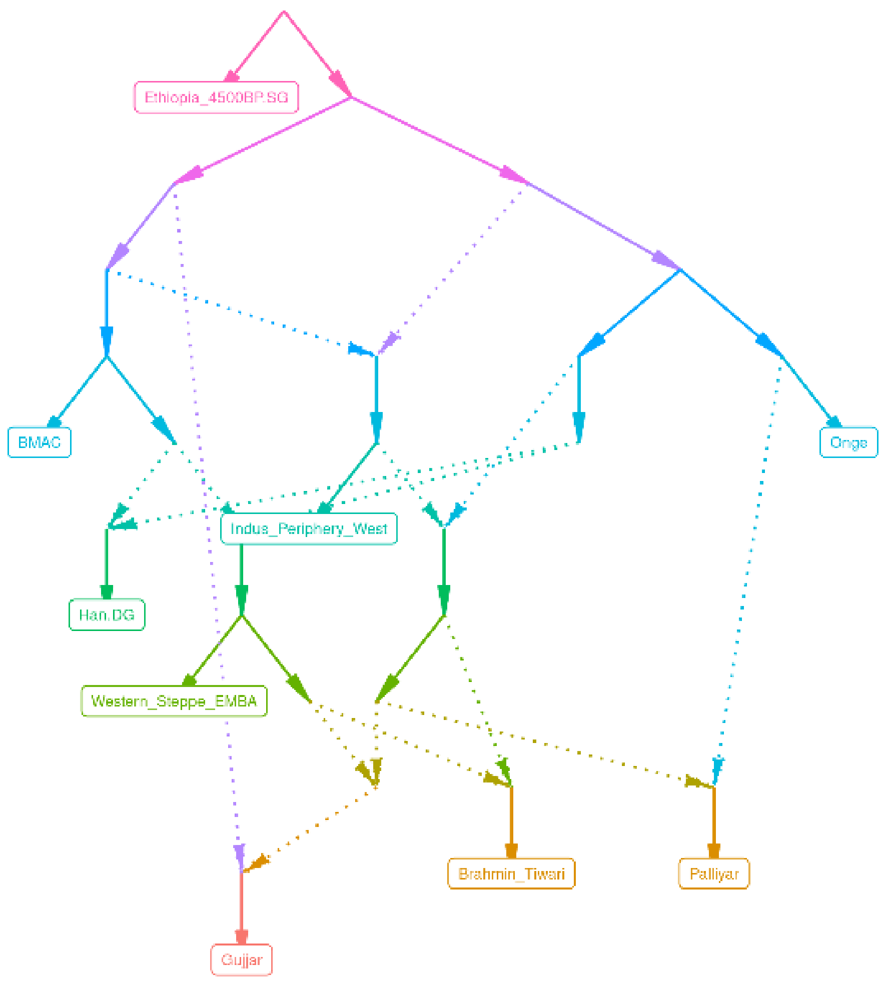

**Fig S6. Geographical population structure inferred from EEMS in the form of effective migration rates. A.** Contour plot for estimates of effective migration rates across the demes. **B.** A plot showing convergence of MCMC chains (1-10) used to infer posterior distribution of parameters. **C.** Plot showing a correlation between observed vs. fitted dissimilarity between demes.

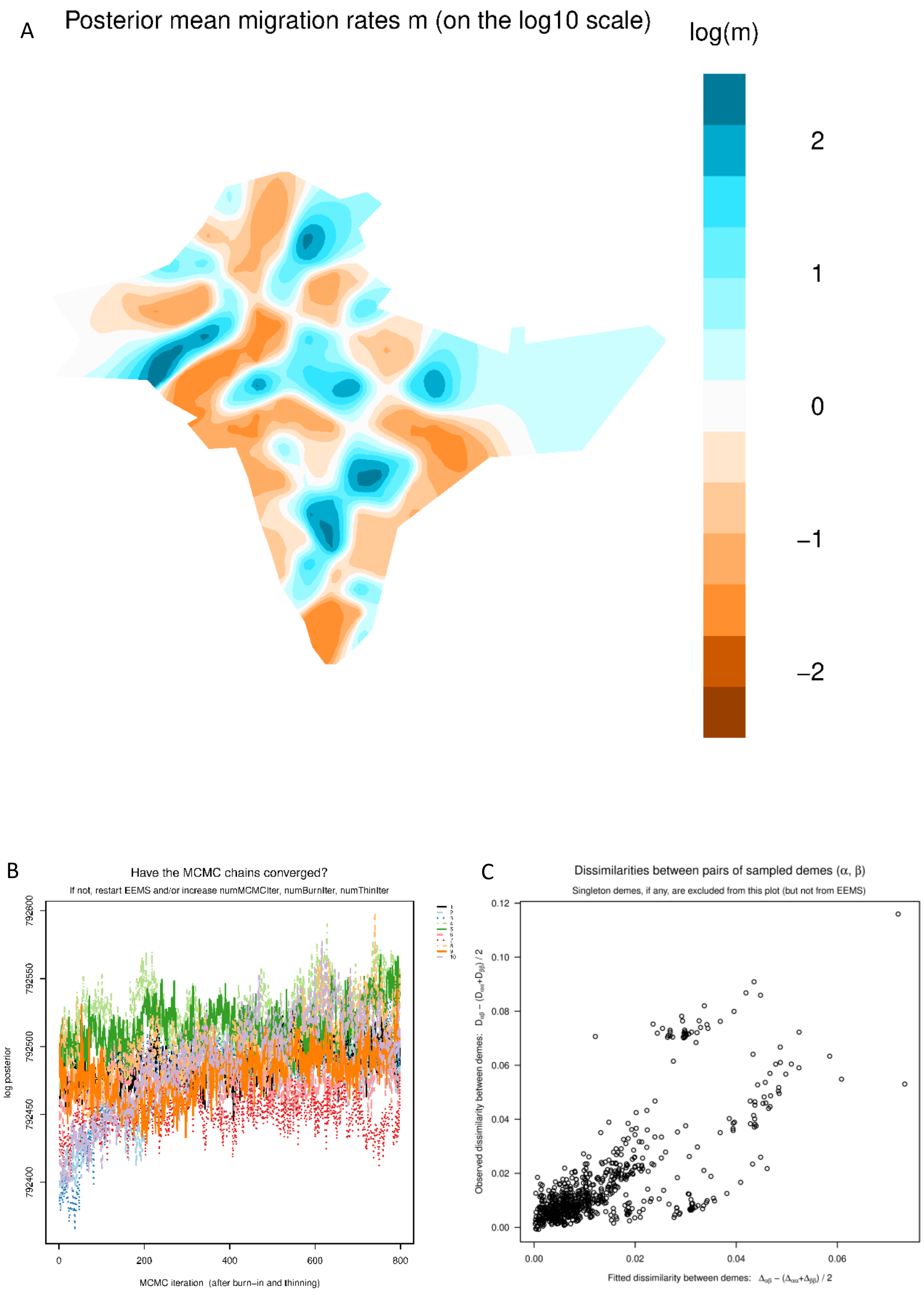

**Fig S7. Population separation history of south west coastal group with other Indian and west Eurasian populations using genome wide genealogy**

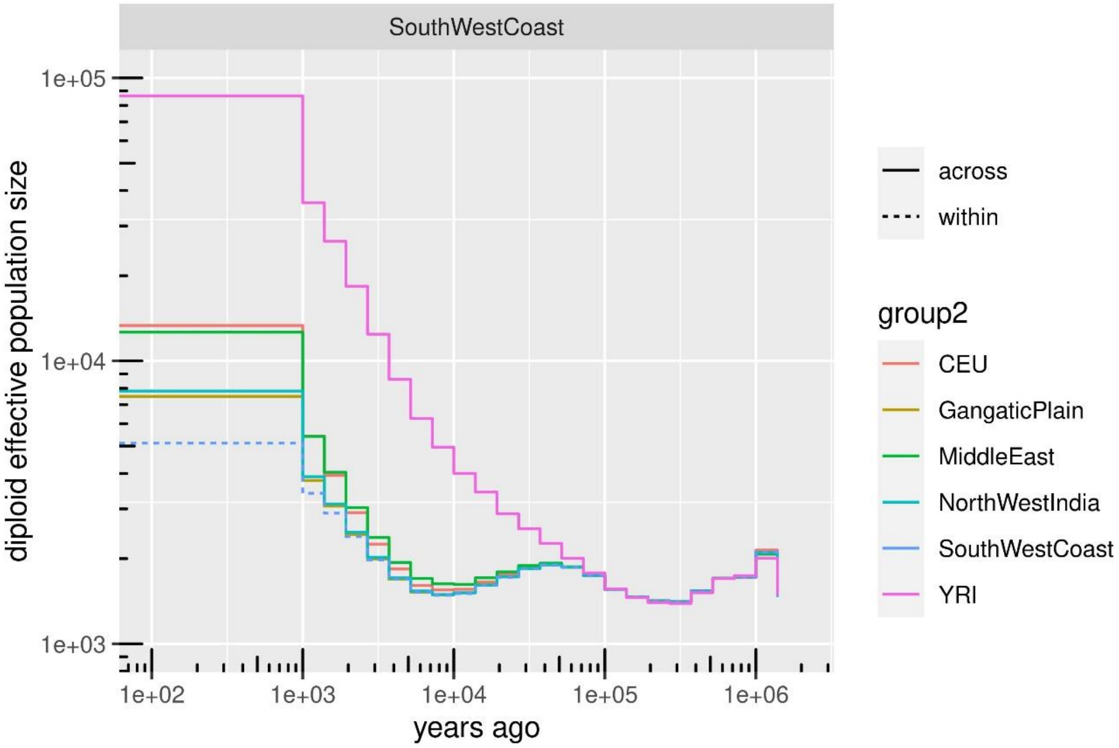

**Fig S8a. Marginal phylogenetic tree around the SNP rs6957904 in the gene *TPK1***

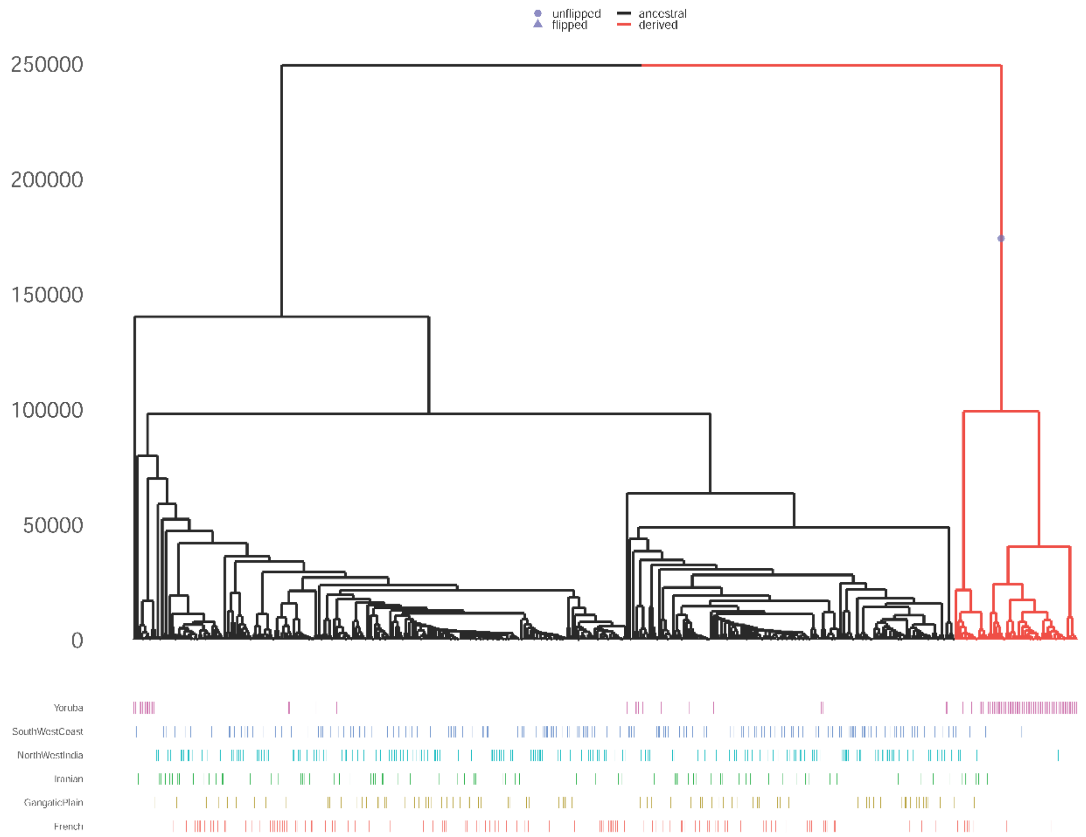


**Fig S8b. Marginal phylogenetic tree around the SNP rs364477 in the gene *DMRT1***

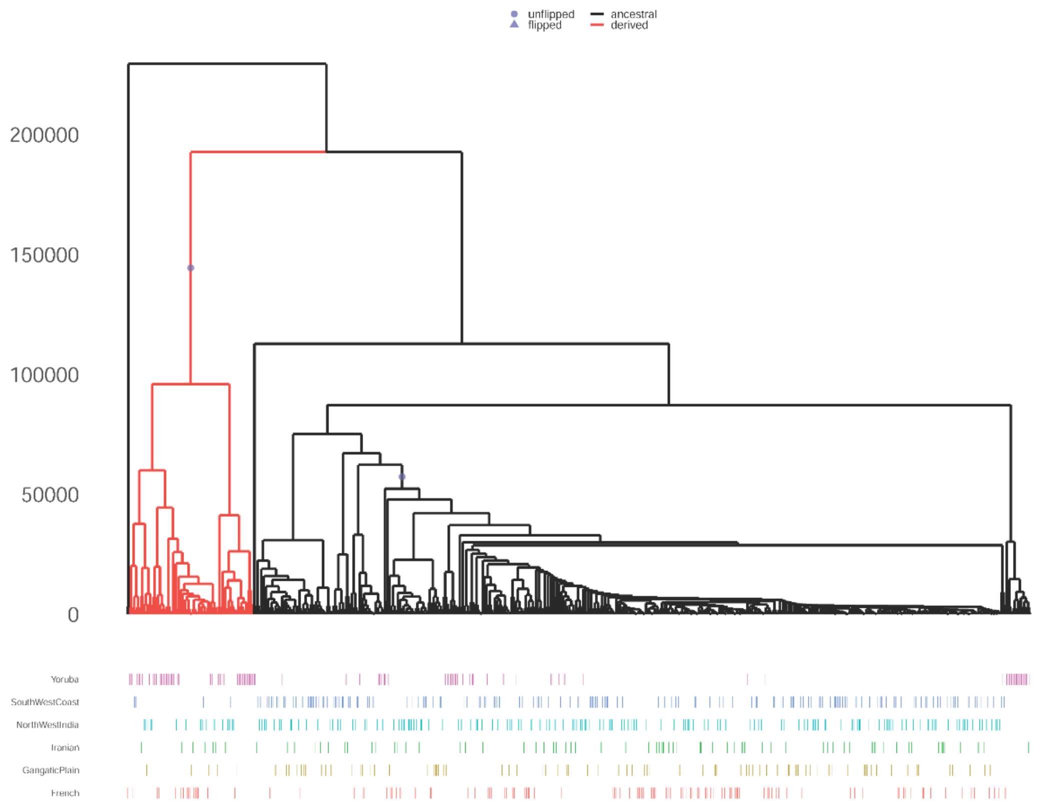

**Fig S8c. Marginal phylogenetic tree around the SNP rs77214450 in the gene *PALLD***

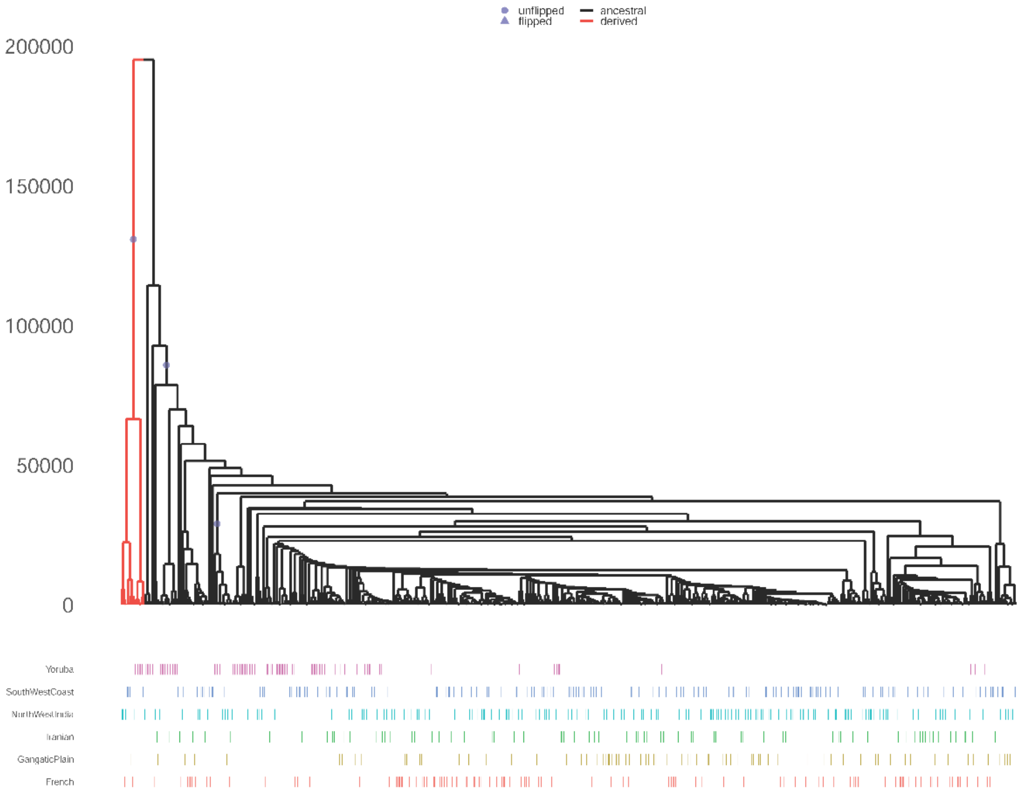


**Fig S8d. Marginal phylogenetic tree around the SNP rs12264666 in the gene *KIAA1217***

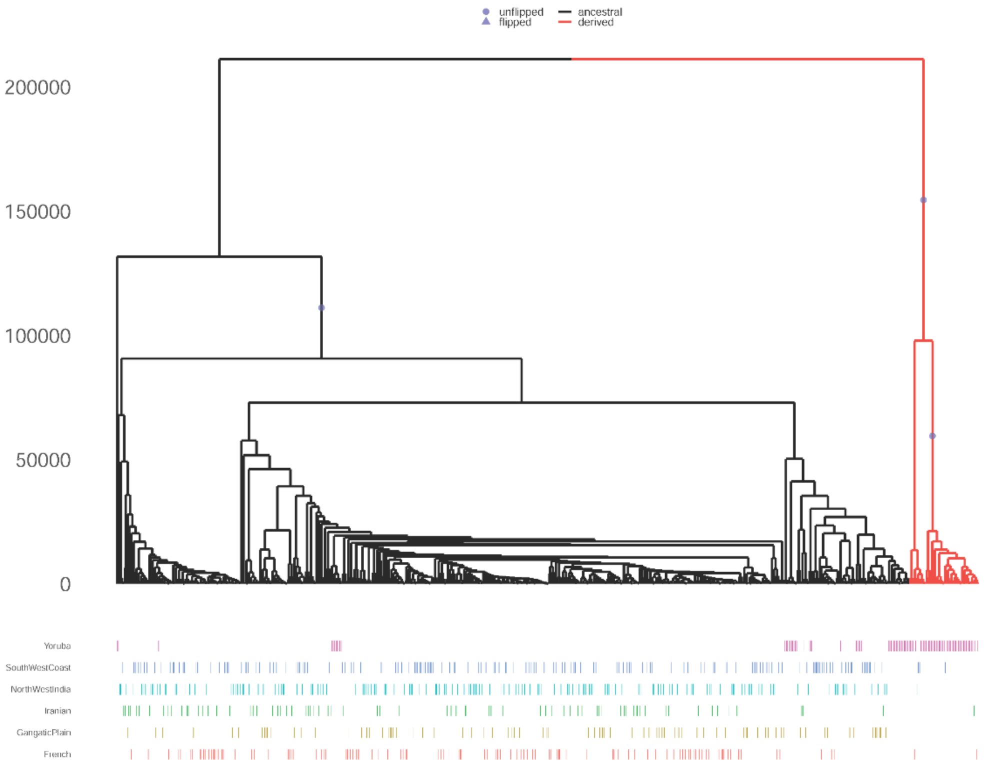

**Fig S9. Mitochondrial haplogroup distribution among three major southwest coastal groups. A.** Distribution of macrohaplogroup among sou west coastal populations. **B.** Distribution of sub haplogroups of mitochondrial haplogroup M. **C.** Distribution of sub haplogroups of mitochondrial haplogroup U. **D.** Distribution of sub haplogroups of mitochondrial haplogroup R.

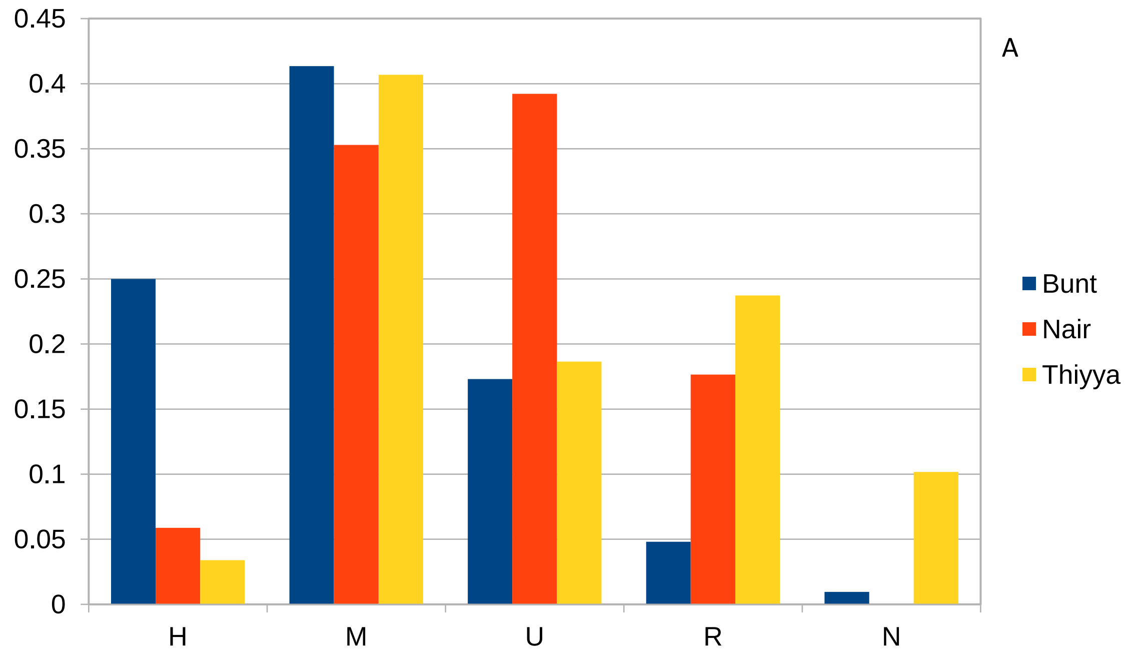


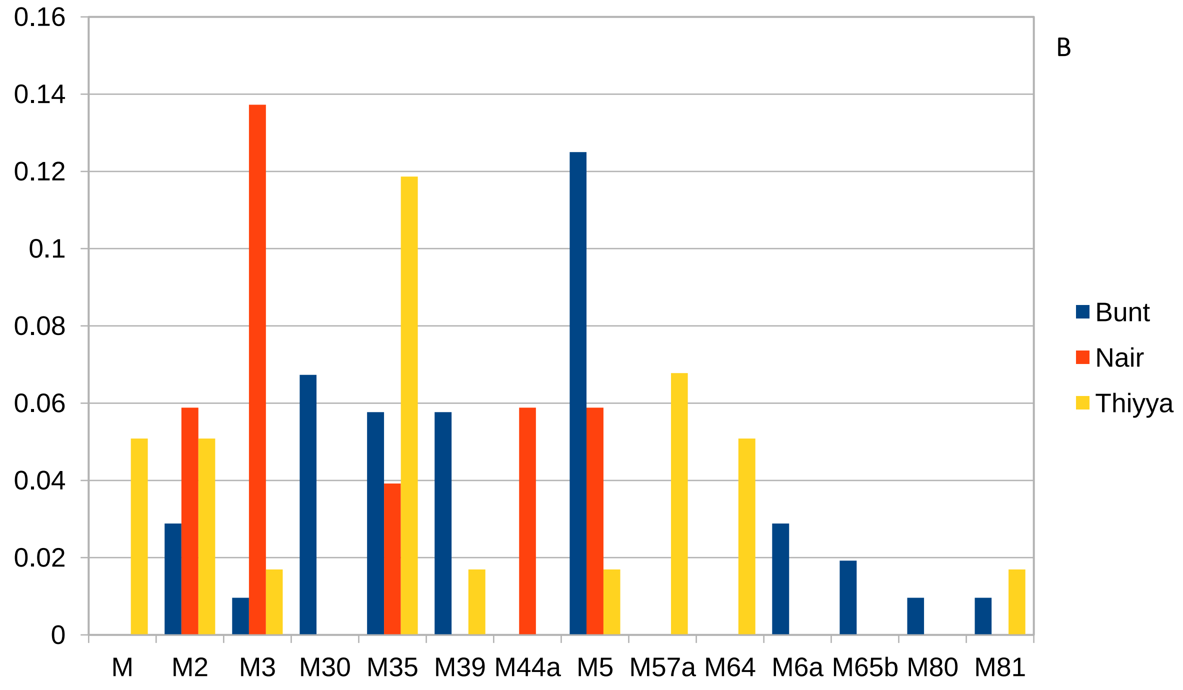

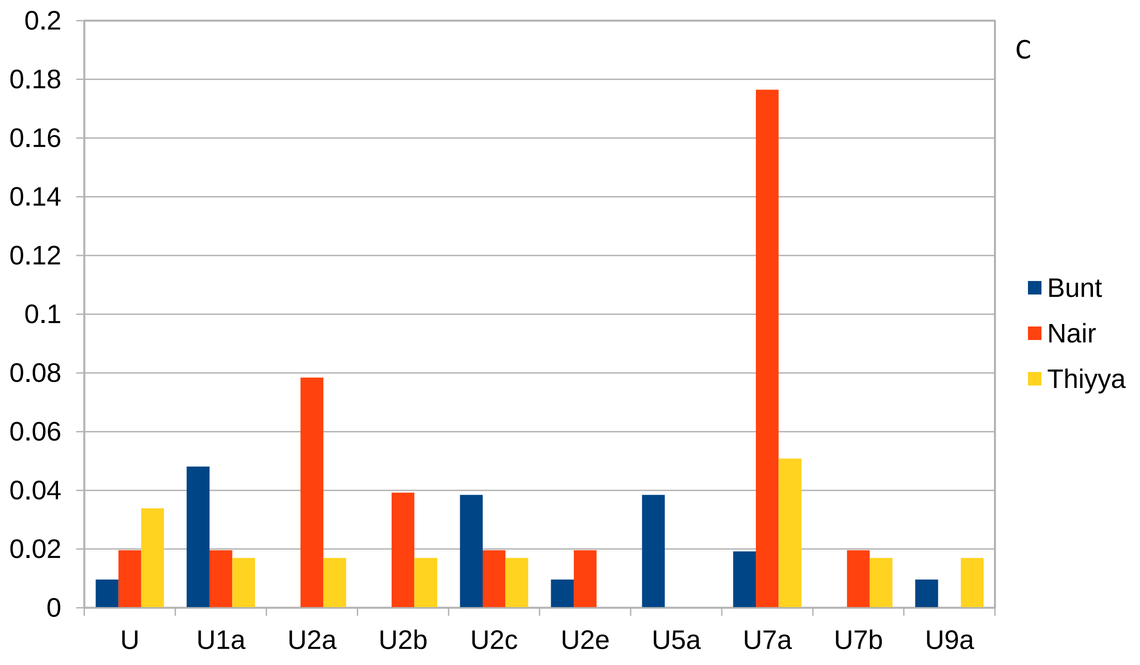
C
0.2
0.18
0.16
0.14
0.12
0.1
0.08
0.06
0.04
0.02
0
U
U1a
U2a
U2b
U2c
U2e
U5a
U7a
U7b
U9a
Bunt
Nair
Thiyya

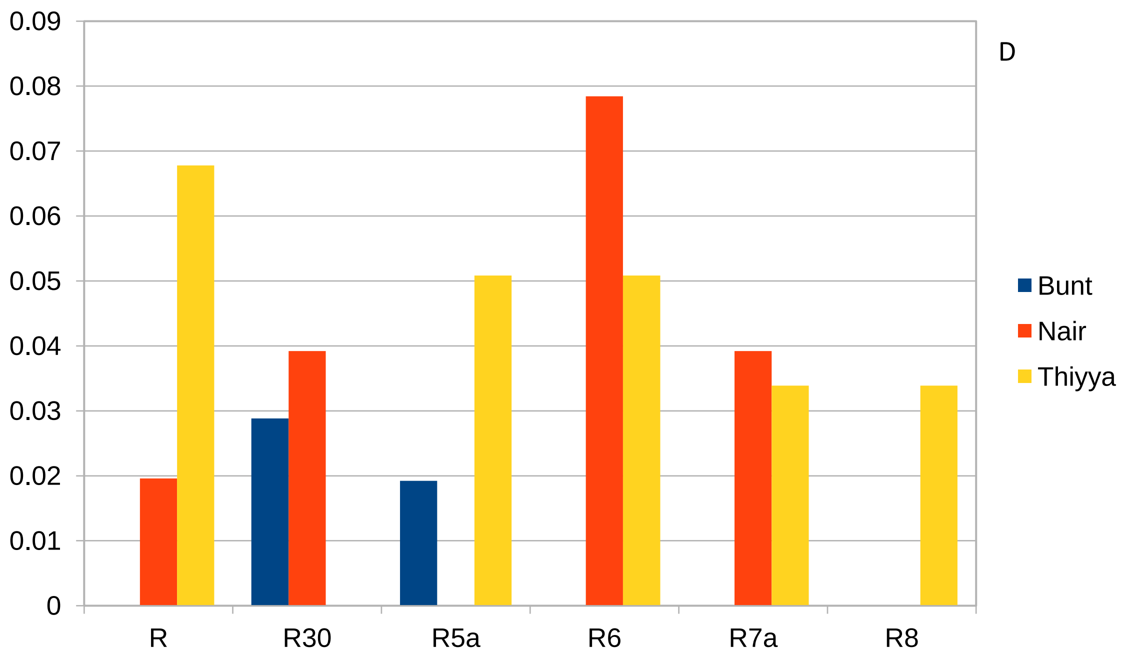
D
0.09
0.08
0.07
0.06
0.05
0.04
0.03
0.02
0.01
0
R
R30
R5a
R6
R7a
R8
Bunt
Nair
Thiyya
